# Lymphatic egress recycles tumor-experienced effector CD8 T cells to sustain immune surveillance

**DOI:** 10.64898/2026.04.30.721705

**Authors:** Ines Delclaux, Katherine S. Ventre, Naomi R. Besson, Tara Muijlwijk, Milad Ibrahim, Guanning Wang, Taylor A. Heim, Maria M. Steele, Alexander C. Huang, Markus Schober, Iman Osman, Amanda W. Lund

**Affiliations:** Ronald O. Perelman Department of Dermatology, NYU Grossman School of Medicine, NYU Langone Health, New York, NY, USA; Division of Hematology/Oncology, Department of Medicine, Perelman School of Medicine, University of Pennsylvania, Philadelphia, PA, USA; Tara Miller Melanoma Center, Abramson Cancer Center, University of Pennsylvania, Philadelphia, PA, USA; Abramson Cancer Center, Perelman School of Medicine, University of Pennsylvania, Philadelphia, PA, USA; Institute for Immunology and Immune Health, Philadelphia, PA, USA; Parker Institute for Cancer Immunotherapy, University of Pennsylvania, Philadelphia, PA, USA; Department of Cancer Biology, Perelman School of Medicine, University of Pennsylvania, Philadelphia, PA, USA; Department of Pathology, NYU Grossman School of Medicine, NYU Langone Health, New York, NY, USA; Laura and Isaac Perlmutter Cancer Center, New York University Langone Health, New York, NY, USA; Translational Immunology Center, NYU Langone Health, New York, NY, USA

**Keywords:** lymphatic transport, stem-like T cells, effector T cells, immune checkpoint blockade, melanoma, lymph node metastasis, T cell migration, tumor-draining lymph node

## Abstract

Successful anti-tumor immune surveillance depends on stem-like CD8^+^ T cells that are enriched in tumor-draining lymph nodes (LN), but how they are maintained over time remains poorly understood. Here, we identify a continuous lymphatic circuit that sustains stem-like CD8^+^ T cells. Using photoconversion to fate-map intratumoral T cells we demonstrate that effector cells exit the tumor microenvironment and migrate back to the draining LN. These tumor-specific, migratory effector T cells avoid chronic antigen stimulation, re-express the transcription factor associated with self-renewal, TCF1, and enter a stem-like state in the LN. Antigen presentation in LNs by dendritic cells drives their proliferation thereby inflating the LN stem-like population. Consequently, maintenance of stem-like T cells and ICB response depends on constitutive lymphatic transport, while LN metastasis compromises the stem-like niche, diminishing ICB response. We, therefore, define a continuous, peripheral lymphatic circuit that recycles tumor-experienced effector T cells to fuel durable, systemic immune surveillance.

## Introduction

Cytotoxic CD8^+^ T cells that accumulate in tumors are often inhibited by chronic antigen, which induces terminal differentiation and functional exhaustion^1,2^. Ongoing anti-tumor immune surveillance, therefore, requires continuous sourcing of effector T cells to the tumor microenvironment, which depends upon a population of stem-like T cells^3–5^ that are enriched in tumor-draining lymph nodes (LN) in both preclinical models^4–8^ and patients^4–6,9,10^. Consistent with a model where tumor-draining LNs are a key source for anti-tumor effectors, systemic rather than intratumoral reinvigoration is necessary for response to immune checkpoint blockade (ICB) and clinical studies associate the abundance of stem-like T cells in the tumor-draining LN with patient response. Therefore, preclinical and clinical data point to an anatomically distant T cell pool that refuels the intratumoral response over time. While recent work has revealed a requirement for dendritic cells (DC) and antigen in maintaining stem-like T cells in the LN^8,11,12^, how this pool of stem-like T cells is chronically maintained and if it is disrupted by standard of care therapy and regional tumor progression^13,14^ remains largely unknown.

Stem-like T cells in the LN are required to expand a circulating early effector population that migrates to the tumor^4,5,8^. Emerging evidence also indicates that migratory capacity may be directly linked to T cell state, with more migratory cells capable of escaping chronic antigen-induced dysfunction in chronic infection and cancer. The transcription factor, Krüppel-like factor 2 (KLF2) regulates T cell trafficking^15–18^ and when enforced, shifts the balance between effector and exhausted lineages^19–21^, likely by maintaining a stem-like population and allowing escape from sites of high antigen burden. While these studies propose a causal link between T cell migration, maintenance of stem-like T cells, and a more functional and persistent effector response, the exact migratory path taken by circulating T cells remains unresolved, limited in part by using static time points from limited tissues to then infer and reconstruct dynamic systemic immune responses.

Importantly, effector and memory T cell migration is not unidirectional, with peripheral tissue exit (egress) via lymphatic vessels back to draining LNs evident across tissues and pathological contexts^22–28^. We recently demonstrated that while tumors retain dysfunctional, exhausted T cells, a broad repertoire of functional, antigen-specific T cells egress the tumor and recirculate to tumor-draining LNs^24^. Inhibiting T cell egress leads to short-term retention and benefits acute response to ICB by sequestering effectors within the tumor parenchyma. However, in line with the impact of KLF2, enforcing long-term residence may drive dysfunction and therefore, migration of effector T cells out of the tumor could be hypothesized to provide a way to escape chronic antigen and the dysfunctional tumor microenvironment. Whether the migration of T cells out of the tumor provides an anatomic route to maintain stemness and effector anti-tumor responses, however, has not been explored.

In this study, we find that the recirculation of tumor-specific T cells from the tumor back to the tumor-draining LN continuously seeds an antigen-dependent stem-like niche, thereby fueling persistent, systemic immune surveillance and response to ICB. This circuit, which is fueled both by antigen and effector T cell recirculation, is disrupted by surgical resection and LN metastasis, indicating that functional lymphatic transport and preservation of the tumor-draining LN are critical to not only initiate, but to maintain systemic anti-tumor immune surveillance for persistent tumor control.

## Results

### Tumor-specific, effector T cells egress tumors

We recently demonstrated that tumor-specific, functional CD8^+^ T cells egress from the tumor via the tumor-associated lymphatic vasculature^24^. These data highlighted the functionality of this migratory, tumor-specific T cell population but raised the question as to their fate upon exit. Interestingly, CD8^+^ T cells similarly egress out of virally infected skin^23^ and lungs^22^ and consequently seed a specialized population of protective, resident memory T cells within the draining LN that are also found in melanoma patients^29^. To, therefore, understand the fate of tumor-egressing CD8^+^ T cells we mapped T cell state in murine melanomas using the Kaede transgenic (Kaede-tg) photoconvertible mouse model^27–29^. We performed single-cell RNA sequencing (scRNAseq) on intratumoral (CD45^+^)^24^ and LN photoconverted (Kaede-red CD45^+^) cells 24 hours post photo-labeling of YUMMER1.7 melanoma immune infiltrates (21 days post intradermal implantation). CD8^+^ T cells (*Cd8a*) were subset from the integrated data and reclustered. Using uniform manifold approximation and projection (UMAP) to visualize the data, we identified clusters corresponding to previously described T cell states: cluster 0 (memory-like; *Sell, Lef1*), cluster 1 (effector; *Ccl5, GzmA*), cluster 2 (Interferon (IFN)-responsive; *Ifit3, Isg15*), cluster 3 (proliferating; *Mki67, Top2a*), cluster 4 (stem-like; *Slamf6, Tox, Pdcd1, Ccr7*), and cluster 5 (dysfunctional; *Havcr2, Pdcd1*) (**Fig. 1A and Extended Data Fig. 1A-C**). We evaluated the relative contribution of intratumoral and egressed T cells to each of these clusters and observed a distinct repertoire of T cell states in the tumor relative to those that had left. Memory-like and stem-like clusters had predominantly egressed, while IFN-responsive, dysfunctional, and proliferating clusters were enriched in the tumor, and only the effector cluster was equally represented in both locations (**Fig. 1B and Extended Data Fig. 1D**). In line with our prior work, and the role of antigen in promoting intratumoral retention^24,30^, we found an increasing differentiation and T cell receptor (TCR) stimulation gradient from egressed to retained, indicating that less differentiated T cells were more likely to exit the tumor microenvironment (**Extended Data Fig. 1E-H**). Consistently, egressing T cells expressed higher levels of *Klf2* (**Fig. 1C**), a master transcription factor that regulates T cell migration via *S1pr1* and *Sell*^29–32^. These data reveal an association between terminal effector differentiation, dysfunction, and tumor retention, with more functional populations leaving the tumor microenvironment and recirculating to tumor-draining LNs, raising the possibility that the process of migration is critical to maintaining these less differentiated states.

**Figure 1.**
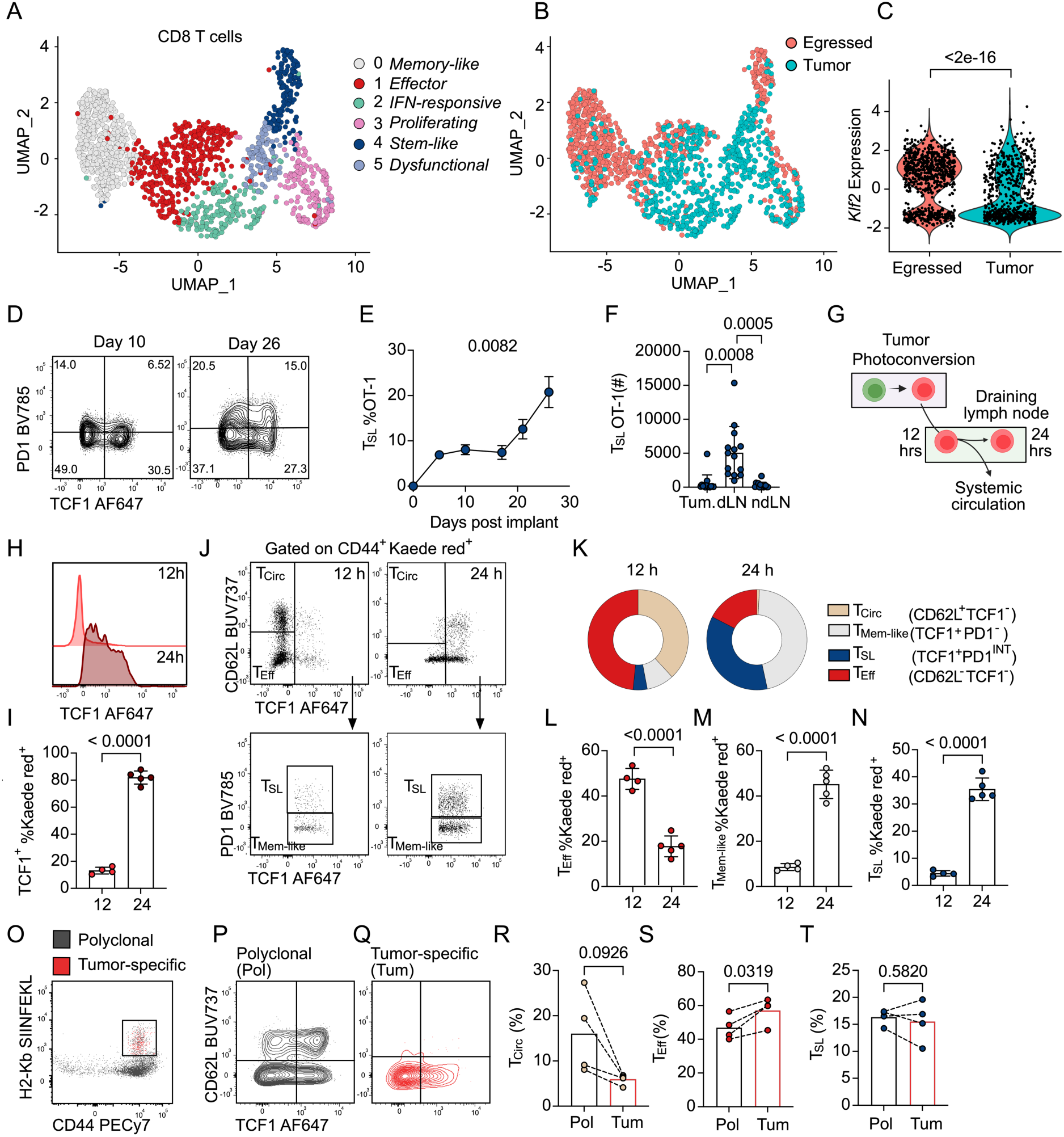
Tumor-specific, effector T cells egress the tumor. **(A)** Uniform Map Approximation Manifold (UMAP) clustering of CD8^+^ T cells sorted from YUMMER1.7 tumors and draining lymph nodes (dLN, Kaede-red^+^ egressed) 24 hours post photoconversion. **(B)** UMAP split by tissue of origin. **(C)** *Klf2* expression in egressed or tumor T cells. **(D)** Representative plots of TCF1 and PD1 gated on CD44^+^ OT-1 T cells in the dLN at day 10 and 26 post YummOVA implantation. **(E)** Percent of stem-like T cells (TCF1^+^PD1^INT^, T_SL_) of CD44^+^ OT-1 T cells over time. **(F)** Number of T_SL_ OT-1 T cells in the tumor, dLN and non-dLN (ndLN) 26 days post YummOVA implantation. **(G)** Diagram of Kaede labeling experiment at 12 and 24 hours. **(H).** Representative histogram of TCF1 and **(I)** percent of TCF1^+^ cells of Kaede-red^+^CD44^+^CD8^+^ T cells in YUMMER1.7 dLN 12 and 24 hours after photoconversion. **(J)** Representative gating strategy of CD44^+^ Kaede-red^+^ CD8 T cells 12 hours and 24 hours post-photoconversion. **(K)** Proportion of T cell states 12 and 24 hours post-photoconversion. **(L).** Percent of effector T cells (CD62L^-^TCF1^-^, T_Eff_), **(M)** memory-like (TCF1^+^PD1^-^, T_Mem-like_), and **(N)** stem-like (TCF1^+^PD1^INT^, T_SL_) T cells of Kaede-red^+^ cells in YUMMER1.7 dLN 12 and 24 hours post-photoconversion. **(O)** Representative plots of CD44 and H2-K^b^-SIINFEKL tetramer staining of Kaede-red T cells in dLN 24 hours post photoconversion of YummOVA tumors. **(P)** Representative plots of CD62L and TCF1 expression in polyclonal (CD44^+^, grey) and **(Q)** tumor-specific (CD44^+^H2-K^b^-SIINFEKL-specific, red) Kaede-red^+^ T cells. **(R)** Percent of circulating (CD62L^+^, T_Circ_), **(S)** T_Eff,_ and **(T)** T_SL_ in polyclonal (Pol) and tumor-specific (Tet) cells. For (**C**) data was analyzed using a two-sided Wilcoxon rank-sum test. For (**E**), each dot is the mean and error bars represent the SEM, data was analyzed using one-way ANOVA adjusted for multiple comparisons (d5, n= 10, d10, n=10, d17 n=7, d21 n=8, d26 n=14). For (**F-R**) each dot represents a mouse. Data was analyzed using one-way ANOVA adjusted for multiple comparisons (**F**), two-sided, unpaired Student’s t-test **(I, L, M, N**), and paired Student’s t-test (**R, S, T**).

We were intrigued by the photo-labeling of a population of LN stem-like T cells given their important role in ongoing tumor control and response to ICB. Stem-like T cells (T_SL_), which we define here as CD8^+^CD44^+^TCF1^+^PD1^INT^, are expanded in the setting of chronic antigen stimulation, and are enriched in tumor-draining LNs in mouse and human^4,5,8,10^. Indeed, we find that tumor-specific stem-like T cells in YummOVA bearing mice, both OT-1 TCR-tg and endogenous (H2K^b^-SIINFEKL; **Extended Data 1I-J**), selectively accumulate in the tumor-draining LN **(Fig. 1D and E)** relative to both their primary tumors and contralateral non-draining LN controls **(Fig. 1F)**, a finding which is consistent across multiple melanoma models (**Extended Data Fig. 1K**). It was therefore surprising that our photoconversion experiments labeled the stem-like population in the LN as it was not evident within the tumor itself. While possible that a rare population of stem-like T cells in the tumor preferentially egress relative to other populations, there was also the possibility that we were capturing a stem-like state that did not migrate out of the tumor at all but rather was induced upon reentry to the LN.

While we and others have primarily analyzed egressing cells at 24 hours^24,31^, the first DCs reach the LN from skin in as early as 8 hrs^32^. Consistently, we noted that there was a significantly higher rate of T cell egress just 12 hours post labeling relative to 24 hours (**Extended Data Fig. 1L**). We, therefore, used these two time points post photoconversion to timestamp intratumoral T cells (day 21 YUMMER1.7) and identify the states most likely to first exit the tumor microenvironment (12 hours) and their subsequent accumulation and fate in the LN (24 hours) (**Fig. 1G**). As seen previously^25^, egressed T cells 24 hours post photoconversion exhibit high expression of TCF1 but, surprisingly, the cells that first leave the tumor (12 hours) did not (**Fig. 1H and I**). Given that all T cells were labeled at the same time, these data were consistent with a shift in T cell state upon entry into the LN. In agreement with our transcriptional analyses, we defined four labeled endogenous CD8^+^CD44^+^ T cell subsets at the two time points: circulating (CD62L^+^TCF1^-^, T_CIRC_), effector (CD62L^-^TCF1^-^, T_EFF_), a memory-like population that had variable expression of CD62L and expressed TCF1 (CD62L^+/-^TCF1^+^PD1^-^,T_mem-like_), and stem-like (CD62L^-^TCF1^+^PD1^INT^, T_SL_) (**Fig. 1J**). Based on this sub-setting strategy, we found that in the first wave of migration (12 hrs) the majority of cells were either circulating or effector T cells (**Fig. 1J-N**). Twelve hours later (24 hrs), there was a drop in CD62L expression on all Kaede-red^+^ T cells and a specific loss of the circulating population (**Extended Data Fig. 1M**-**O**). Effector T cells were also reduced (**Fig. 1L**), while memory-like and stem-like cells enriched in the LN over time (**Fig. 1 M-N**). Circulating T cells were more prevalent in the endogenous polyclonal pool than in the tumor-specific (H2-K^b^-SIINFEKL, **Fig. 1O-R**) population, likely indicating that they represent a bystander population that trafficked from the tumor and passed through the first draining LN within the 24 hours post labeling. In contrast, effector and stem-like T cells were equally represented in both polyclonal and tumor-specific populations (**Fig. 1S-T**), suggesting that these cells were tumor-specific rather than bystanders. This time-stamped labeling strategy, therefore, revealed a functional distinction between the T cell states actively exiting the tumor microenvironment from those that accumulate over time within the draining LN. This data required a reinterpretation of our original scRNAseq, which represented the fate of T cells labeled in the tumor 24 hrs prior and suggested that egress is associated with the re-activation of TCF1 and the emergence of memory-like and stem-like T cells within the LN.

### Tumor-egressing effector T cells seed LN stem-like T cells

Based on these photo-labeling experiments, we postulated that tumor-specific T cells egress the tumor via lymphatic vessels, traffic to the LN, and convert to stem-like T cells, which progressively accumulate over time (**Fig. 1E**). To determine the migratory requirements to maintain the LN stem-like T cell population, we first used αCD62L to block hematogenous migration into the LN. Inhibition of LN influx did not reduce the number of stem-like T cells (**Extended Data Fig. 2A**) despite the expected significant reduction in naive CD8^+^ T cells (**Extended Data Fig. 2B**). We further observed enrichment of stem-like T cells within activated endogenous (CD44^+^) (**Fig. 2A**) and tumor-specific TCR-tg T cells (**Fig. 2B**), in line with the reduction in blood-derived naive and memory T cells. Similar results were observed in a second melanoma model, B16OVA (**Extended Data Fig. 2C**). Therefore, the stem-like T cell population does not depend upon continuous seeding from the blood, suggesting either a resident phenotype or an alternative route of trafficking, with our photo-labeling experiments supporting at least a contribution from the latter.

**Figure 2.**
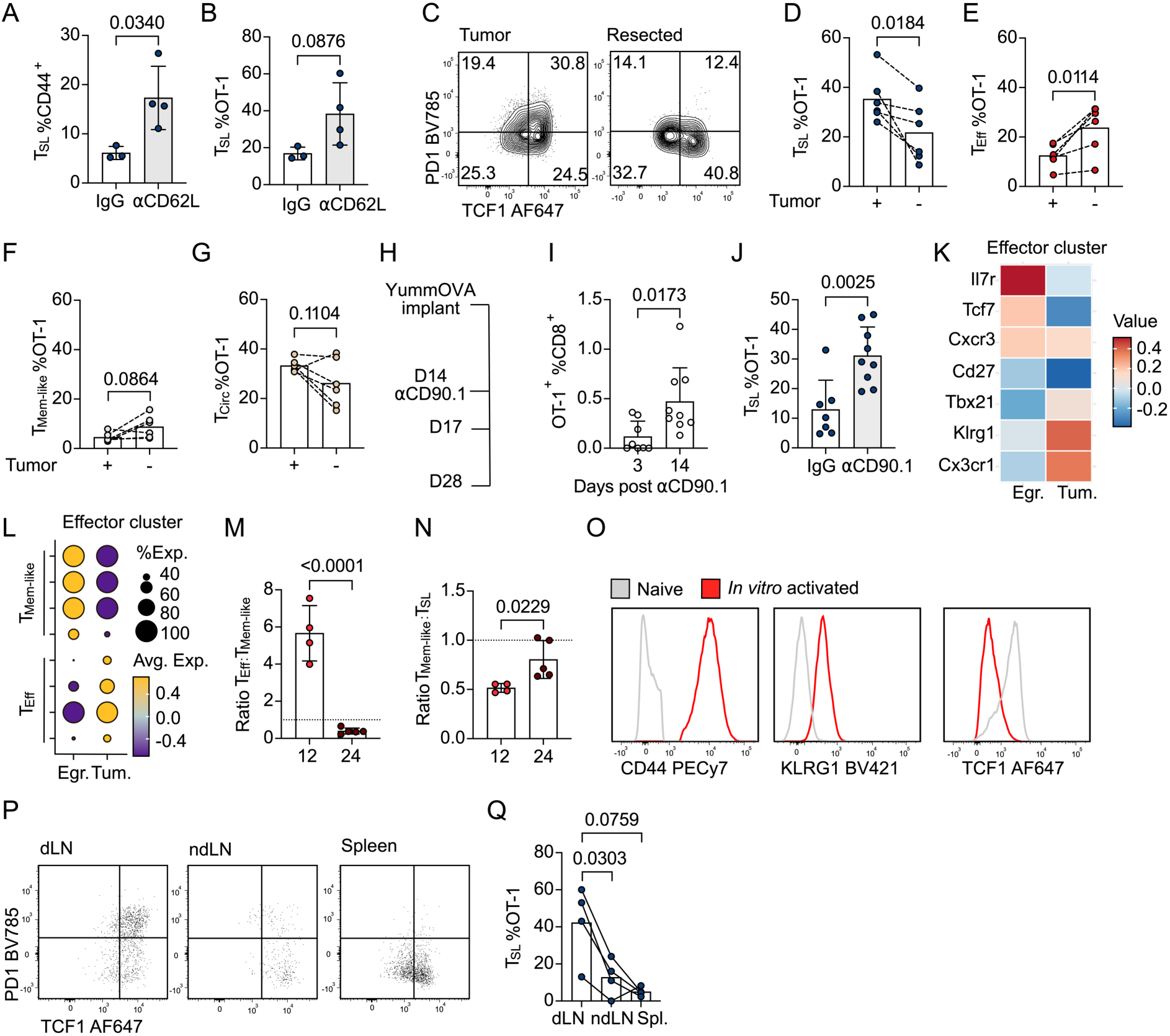
Egressing early effector T cells seed LN stem-like T cells. **(A)** Percent of stem-like (TCF1^+^PD1^INT^, T_SL_) of CD44^+^ CD8 T cells, and **(B)** percent of T_SL_ of CD44^+^ OT-1 T cells in YummOVA draining lymph nodes (dLN) treated with IgG or αCD62L. **(C)** Representative plots showing TCF1 and PD1 expression gated on CD44^+^ OT-1 T cells in LNs draining intact (Tumor) or resected (Resected) YummOVA tumors. **(D)** Percent of T_SL_ **(E)** Effector, (CD62L^-^TCF1^-^PD1^-^, T_Eff_), **(F)** memory-like, (CD62L^-^TCF1^+^PD1^-^, T_Mem-like_), and **(G)**. circulating (CD62L^+^, T_Circ_) of CD44^+^OT-1 T cells in the LN draining intact (+) or resected (-) tumor. **(H)** Schematic of depletion experiment timeline. **(I)**. Percent of OT-1 of CD8^+^ T cells in the dLN 3- or 14-days post αCD90.1 **(J)** Percent of T_SL_ of CD44^+^OT-1 T cells in the dLN 14 days after IgG or αCD90.1 treatment. **(K)** Heatmap showing gene expression in egressed (Egr.) or tumor-retained (Tum.) effector cluster. **(L)** Memory-like or Effector signature scores^19,33–35^ in Egr. or Tum. effector cluster **(M)** Ratio T_Eff_ to T_Mem-like_ and **(N)** T_Mem-like_ to T_SL_ Kaede-red^+^ T cells in the dLN 12 or 24 hours post-photoconversion of YUMMER1.7 tumors. **(O)** Representative histograms of indicated markers in naïve OT-1 (grey) or *in vitro* activated OT-1 T cells (red). **(P)** Representative plots of TCF1 and PD1 expression of transferred OT-1 T cells 4 days after transfer in YummOVA tumors (d.9 post implantation) in the dLN, ndLN and spleen. **(Q)** Percent of T_SL_ of CD44^+^OT-1 T cells in the dLN, dLN and spleen 4 days after transfer. For all analyses each dot represents a mouse. Data was analyzed, two-sided, unpaired Student’s t-test **(A, B, I, J, M, N**), and paired Student’s t-test (**D, E, F, G**), and one-way ANOVA adjusted for multiple comparisons (**Q**).

To directly test the necessity of lymphatic transport from tumors to accumulate stem-like T in the LN, we transferred OTI TCR-tg T cells into mice and co-implanted YummOVA tumors intradermally so that each drained to a brachial LN. Ten days after tumor implantation, following initial priming and when we just begin to observe the accumulation of stem-like T cells (**Fig 1E**), we resected one tumor preventing transit through the primary tumor while leaving the other tumor and systemic anti-immune response intact^23^. Analysis of the LNs 27 days post-implantation (16 days post-resection), revealed that the LN draining the resected tumor had significantly fewer stem-like T cells (**Fig. 2C and D**) and a larger effector population compared to the contralateral LN with an intact tumor (**Fig. 2E**). Memory-like T cells also showed a slight bias toward the resected LN, whereas circulating T cells equally distributed between both LNs (**Fig. 2F and G**).

These data indicated that the expansion of the stem-like pool over time depends on constitutive lymphatic transport and cannot be maintained by a LN resident population that emerges upon priming. To specifically determine if T cells that migrate through the afferent lymphatic vessels seed the stem-like population in the tumor-draining LN, we transferred congenically marked (CD90.1), naive OT1 TCR-tg T cells prior to YummOVA implantation. At day 14 post-tumor implantation, we treated mice with αCD90.1 antibody (**Fig. 2H**) effectively depleting all OT-1 T cells in blood (**Extended Data Fig. 2E**) and LNs (**Fig. 2I and Extended Data Fig. 2F-H**), while maintaining T cells in the tumor (**Extended Data Fig. 2F and 2I**)^23^. We leveraged this system to lineage trace tumor-infiltrating T cells and map their accumulation in different tissues over time. Fourteen days post depletion, we found that CD90.1^+^OT-1 T cells had preferentially re-populated the tumor-draining LN (**Fig. 2I**) relative to non-draining, contralateral LN controls (**Extended Data Fig. 2J**), and the spleen (**Extended Data Fig. 2K**). In addition to the numerical rebound of the OT-1 population in the tumor-draining LN, we observed enrichment for stem-like T cells relative to un-depleted controls (**Fig. 2J**), together showing that tumor-egressing T cells are sufficient to repopulate stem-like T cells in tumor-draining LNs. Of note, in contrast to what we had seen following skin infection^23^, resident memory-like T cell cells (CD62L^-^CD69^+^CXCR6^+^, TRM) were not efficiently repopulated by tumor-egressing T cells over this time frame (**Extended Data Fig. 2L**).

Our photolabeling experiments indicated that the cells most likely to migrate out of the tumor were in a circulating or effector state. Given that the circulating phenotype seemed to match to a likely bystander population (**Fig. 1P-R**), we predicted that the tumor-egressing effector T cell was the precursor to the LN stem-like state. Upon re-examination of the effector cluster within our transcriptional dataset (**Fig. 1A**), we noted that the tumor retained and egressed effector populations differentially expressed genes associated with migration and effector function. Transcripts associated with migration and a memory precursor state (*Il7r, Tcf7, Cxcr3*, *Cd27*) where enriched in the tumor-egressed fraction of the effector cluster, whereas the intratumoral effector T cells expressed higher levels of *Tbx21, Klrg1, Cx3cr1*, genes associated with the effector state (**Fig. 2K**). Accordingly, egressed T cells from the effector cluster scored higher for memory precursor signatures while retained T cells scored for effector function (**Fig. 2L**)^19,33–35^. When looking at the cell populations, we also observed a shift in cell populations over time: the effector-to-memory-like ratio flipped between 12 and 24 hours post-photoconversion (**Fig. 2M**), followed by a subsequent shift from memory-like to stem-like (**Fig. 2N**), altogether consistent with a state conversion upon tumor egress.

Early effector T cells may regain memory potential through a reversal of the epigenetic silencing of *Tcf7*, which allows memory to be tuned to the peripheral effector response^36^. We, therefore, envisioned a model wherein an effector T cell that escapes chronic antigen engagement in the tumor microenvironment^24^ can subsequently activate memory and stem-like programs in the tumor-draining LN. To test this specifically, we asked if TCF1^-^ effector cells could reestablish TCF1 expression and engage a stem-like state in the tumor-draining LN following tumor exit. To do this, we activated OT-1 T cells *in vitro* and transferred them intravenous into YummOVA tumor-bearing mice at day 9 post-implantation. In vitro activated T cells present an effector-like phenotype, upregulating activation markers such as CD44, KLRG1 (**Fig. 2O**), PD1, CD69 (**Extended Data Fig. 2M**) and downregulating CD62L and TCF1 compared to naive T cells (**Fig. 2O and Extended Data Fig. 2M**). CD62L antibodies were delivered together with the T cell transfer to prevent the migration of in vitro activated T cells into LNs through high endothelial venules, requiring tumor transit to access the draining LN. We found that the transferred T cells readily populated the tumor-draining LN, spleen, and to a lesser extent the non-draining LN 4 days post transfer (**Fig. 2P**). In line with our proposed model, the vast majority of the transferred T cells had reactivated TCF1 expression in all compartments (**Fig. 2P**), however, only in the tumor-draining LN did we find TCF1^+^PD-1^INT^ stem-like T cells (**Fig. 2Q**). Together, these results indicate that T cell tumor egress is sufficient to populate the stem-like pool in the tumor-draining LN and further suggests that a migratory effector T cell retains phenotypic plasticity allowing for reactivation of a memory- and stem-like programs in the tumor-draining LN.

### Chronic antigen in the LN converts tumor-egressing effectors to stem-like T cells

Our data indicate that TCF1 and a memory-like program is reactivated upon tumor egress but suggested that the nature of the LN environment was critical to induce the stem-like state. Our trafficking experiments further demonstrate at least a transient residence of stem-like T cells, evidenced by their enrichment upon αCD62L blockade and expression of residence-associated genes (*Cxcr6, Bhlhe40*) (**Extended. Data Fig. 1C**), both of which are in agreement with data indicating TGFβ−dependent retention^3,7^. Crucially, antigen encounter can serve as a retention cue both during priming^37^, and in peripheral, non-lymphoid environments^24,38^. In fact, antigen engagement in LNs can promote T cell retention for up to 12 hours^37^, a time frame that matches our experimental design. Given this, despite earlier hypotheses from us and others that T cell egress might protect T cells from antigen encounter, we asked whether antigen re-engagement in the tumor-draining LN was necessary for the acquisition and maintenance of the stem-like T cell state.

In line with a model that antigen engagement in the LN defines the stem-like state, we find that stem-like OT-I TCR-tg T cells were more proliferative than either circulating or effector-like states, which likely contributes to their local enrichment (**Fig. 3A**). This increase in proliferation was associated with chronic antigen engagement (Nur77GFP reporter) in the tumor and tumor-draining LN that persisted well past initial priming (**Fig. 3B-D**), and a relative increase in recent antigen engagement in the stem-like population relative to other OT-1 T cells, despite the fixed TCR (**Fig. 3E**). Cells that had recently engaged antigen (Nur77GFP^+^) were more proliferative than those that had not (**Fig. 3F**), and tumor-egressing T cells upregulated PD-1 upon arrival to the LN (**Fig. 3G**), supporting antigen-dependent induction of the stem-like state by egressing effectors. The pattern of stem-like T cell accumulation also mirrored antigen presence, with T cells preferentially adopting the stem-like state in the ipsilateral brachial and axillary LNs, (**Fig. 3H**) where antigen is present (**Extended Data Fig. 3A**), but not in the spleen (**Fig. 3H**). Instead, the migratory effector state distributes systemically (**Fig. 3I and 3J**).

**Figure 3.**
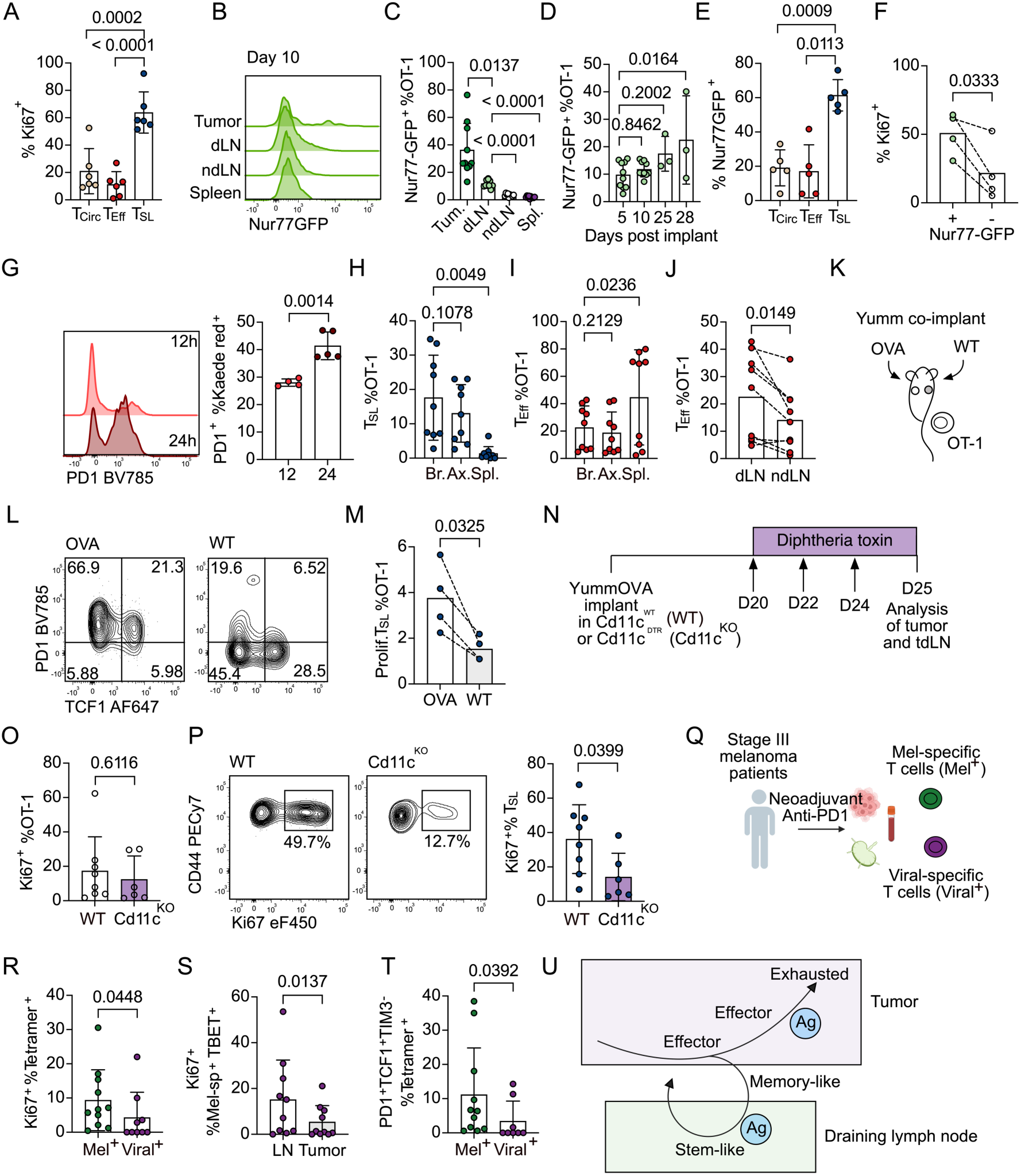
Chronic antigen in the LN converts egressing effectors to stem-like T cells. **(A)** Percent of Ki67^+^ circulating (CD62L^+^, T_Circ_), effector (CD62L^-^TCF1^-^PD1^-^, T_Eff_) and stem-like (TCF1^+^PD1^INT^, T_SL_) T cells in YummOVA draining lymph nodes (dLN) 26-28 days post tumor-implantation. **(B)** Representative histogram of Nur77GFP expression in OT-1 T cells in the tumor, brachial dLN, non-dLN (ndLN) and spleen at day 10 post tumor implantation. **(C)** Percent of Nur77-GFP^+^ of OT-1 T cells in tumor, dLN, ndLN and spleen at day 10 post tumor implantation. **(D)** Percent of Nur77-GFP^+^ of OT-1 T cells in the dLN at the indicated time points post tumor implantation. **(E)** Percent of Nur77-GFP^+^ in the indicated T cell subsets at day 26-28 post tumor implantation. **(F)** Percent of Ki67^+^ in Nur-77GFP^+^ or Nur-77GFP^-^ OT-1 T cells at day 26-28 post tumor implantation. **(G)** Representative histogram (left) and quantification of PD1 expression as a percent of Kaede-red^+^ CD44^+^ CD8^+^ T cells (right) in the dLN at 12 and 24 hours post-photoconversion of YUMMER1.7 tumors. **(H)**. Percent of T_SL_ of OT-1 T cells in brachial (Br.) and axillary (Ax.) ipsilateral LNs and in the spleen 26 days post YummOVA implantation. **(I)** Percent of T_Eff_ of OT-1 T cells in brachial (Br.) and axillary (Ax.) ipsilateral LNs and in the spleen 26 days post YummOVA implantation. **(J)**. Percent of T_Eff_ of OT-1 T cells in ipsilateral brachial dLN and contralateral brachial ndLN. **(K)** Schematic of Yumm/YummOVA co-implantation experiment. **(L)**. Representative plot of TCF1 and PD1 expression of OT-1 T cells in YummOVA (OVA) dLN and Yumm (WT) dLN**. (M)** Percent of Ki67^+^T_SL_ of OT-1 in OVA and WT dLN. **(N)** Schematic showing Cd11c diphtheria toxin experiment. **(O)** Percent of Ki67^+^ of OT-1 in YummOVA dLNs at day 25 post implantation in Cd11c^WT^ (WT) or Cd11c^DTR^ (KO) after 3 diphtheria doses. **(P)** Representative plots (left) and percent (right) of Ki67 expression in T_SL_ OT-1 T cells in Cd11c^WT^ (WT) or Cd11c^DTR^ (KO). **(Q)** Stage III melanoma patients treated with neoadjuvant anti-PD1 and analysis of tumor, LN and blood with tetramer-specific flow cytometry analysis. **(R)** Percent of PD1^+^TCF1^+^TIM3^-^of melanoma-specific or viral-specific T cells in LNs of melanoma patients. **(S)** Percent of Ki67^+^ of TBET^+^ melanoma-specific T cells in LNs or tumors of melanoma patients **(T)** Percent of Ki67^+^ of tetramer^+^ cells in LNs of melanoma patients. **(U)** Proposed model, Ag = Antigen. For **(A-P)**, each dot represents a mouse and for **(R-T)**, each dot represents a patient. Data was analyzed using one-way ANOVA adjusted for multiple comparisons (**A, C, D, E, H, I**), two-sided, unpaired Student’s t-test **(G, O, P**), and paired Student’s t-test (**F, J, M**), Mann-Whitney two-tailed test (**R,S**) and Wilcoxon matched-pairs signed two-tailed rank test (**T**).

Based on these observations, we directly tested the contribution of antigen to the LN stem-like state. First, we co-implanted YummOVA and Yumm tumors on contralateral sides of the same mouse, with distinct draining LNs (**Fig. 3K**). Here a systemic OVA-specific T cell response can transit through both tumors but will only reengage antigen in the LNs draining the OVA-expressing tumor. Analysis of tumor-draining LNs 21 days post implantation revealed an increase in proliferating, stem-like T cells in the OVA-draining LN relative to the contralateral wildtype control (**Fig. 3L and M**), highlighting the necessity of antigen re-engagement in the LN to induce a proliferative stem-like T cell niche.

Since cells engaging antigen adopt the stem-like state, and antigen is necessary to maintain this state in the draining LN, we wondered what role antigen presenting cells in the draining LN, specifically DCs, would have in their maintenance. We implanted YummOVA tumors in Cd11c-DTR mice, where Cd11c^+^ cells express the diphtheria toxin receptor, allowing for their depletion upon diphtheria toxin administration (**Extended Data Fig. 3B**). To allow for initial priming and investigate the impact of DCs on stem-like T cell maintenance rather than formation, we administered diphtheria toxin 20 days post-implantation and analyzed LNs 5 days later (**Fig. 3N**). By day 20 post implantation, we find that the LN stem cell population continues to be further enriched in the antigen-specific OT-I population relative to initial levels formed by day 7 (**Fig 1E**). Diphtheria toxin administration led to a decrease in both MHC-II^+^ Xcr1^+^ cDC1 and MHC-II^+^ Sirpα^+^ Cd11b^+^ cDC2 populations (**Extended Data Fig. 3B-F**) and a significant reduction in total number of OT-1s in the draining LNs (**Extended Data Fig. 3G)**, but no difference in proliferation of the bulk OT-1 T cells (**Fig. 3O**). In contrast, stem-like cells in the draining LNs of Cd11c^DTR^ exhibited a significant decrease in proliferation (**Fig. 3P**), indicating an increased reliance on DCs for their persistence. Taken together, these results show that antigen presentation in the draining LN is required for the long-term maintenance of the stem-like state. This data importantly reinforces the need for continuous DC engagement for maintenance past the initial priming evident in the first week post implantation^11^, and at time points when T cell egress meaningful contributes.

We wondered if the antigen-dependent induction of the stem-like state in LNs is also relevant in LNs of melanoma patients. We re-analyzed data^39^ collected using combinatorial tetramers to identify a pool of melanoma-specific (specific for GP100, AIM2, MART-1, MAGE etc.) and viral-specific (specific for EBV EBNA 3A, CMVpp65, FluNP etc.) T cells in non-metastatic LNs of stage III melanoma patients that had received one dose of anti-PD1 in the neoadjuvant setting (**Fig. 3Q**). In this setting, the abundance of melanoma-specific stem-like CD8^+^ T cells is associated with response to ICB treatment^39^. By directly comparing melanoma and virus-specific CD8^+^ T cells in the same LNs, we found that melanoma-specific T cells were more proliferative than viral-specific T cells in the LN (**Fig. 3R**) and stem-like T cells were also more proliferative in the LN than in the tumor (**Fig. 3S**). Additionally, consistent with our preclinical data, melanoma-specific T cells were more likely to be in the stem-like state than anti-viral T cells in the same LN (**Fig. 3T**), concordant with our findings in mice to indicate that the enrichment of the stem-like state in LNs is proliferation and antigen-dependent.

We therefore propose a model (**Fig. 3U)** in which plastic, effector T cells make a continuous migratory loop from LN to tumor and back to sustain the anti-tumor immune response. Chronic antigen exposure within the tumor microenvironment enforces differentiation into a resident and dysfunctional state. Instead, a subset of these effector T cells egresses the tumor, reactivate expression of TCF-1 and subsequently expand upon antigen re-encounter in the draining LN thereby maintaining the systemic effector response. This ongoing circuit is predicted to depend upon constitutive lymphatic transport that coordinates the concomitant egress of tumor-specific T cells and refuels the antigen niche within the LN.

### Lymphatic transport maintains the antigen-dependent niche and immunotherapy response

The implication of our proposed model (**Fig. 3U**) is that disruption of the lymphatic circuit would gradually deplete the stem-like pool. Given the important role of antigen described above and observed clinical benefit of neoadjuvant over adjuvant therapy, we directly tested how the stem-like pool and response to ICB are affected by either standard of care tumor resection. We first resected YummOVA tumors at day 20 post-implantation analyzing the remaining OT-I TCR-tg CD8^+^ T cells over time (0,7,16, 21 and 30-days post-resection). Lymphatic transport and DC migration is required for antigen transport to draining LNs^40–42^, and consistently, tumor-resection led to a reduction in antigen engagement (Nur77GFP^+^) by OT-1 tumor-specific T cells in the tumor draining LN (**Fig. 4A**). Upon resection, stem-like T cells were progressively lost over time (**Fig. 4B and C**) reaching the background levels observed in non-draining LNs, indicating that the stem-like state requires constitutive lymphatic drainage not only for its early expansion but for long-term maintenance. In contrast, CD62L^+^ circulating T cells were stably maintained over time (**Fig. 4D**), distributed equally between the draining LN and non-draining LN (**Fig. 4E**), and expressed TCF1, signaling a transition to memory. Other T cell states also transitioned to memory upon tumor resection and consequent antigen loss: memory-like T cells (CD62L^-^TCF1^+^PD1^-^) increase in percentage after resection (**Extended Data Fig. 4A**), remaining enriched in the draining LN (**Extended Data Fig. 4B**), resident-like T cells (CD62L^-^CD69^+^CXCR6^+^TCF1^-^) were stably maintained (**Extended Data Fig. 4C**) and, reflecting their resident nature, remain enriched in the draining LN (**Extended Data Fig. 4D**), while an effector memory (CD62L^-^CD69^-^) population distributes systemically (**Extended data Fig. 4E and F**).

**Figure 4.**
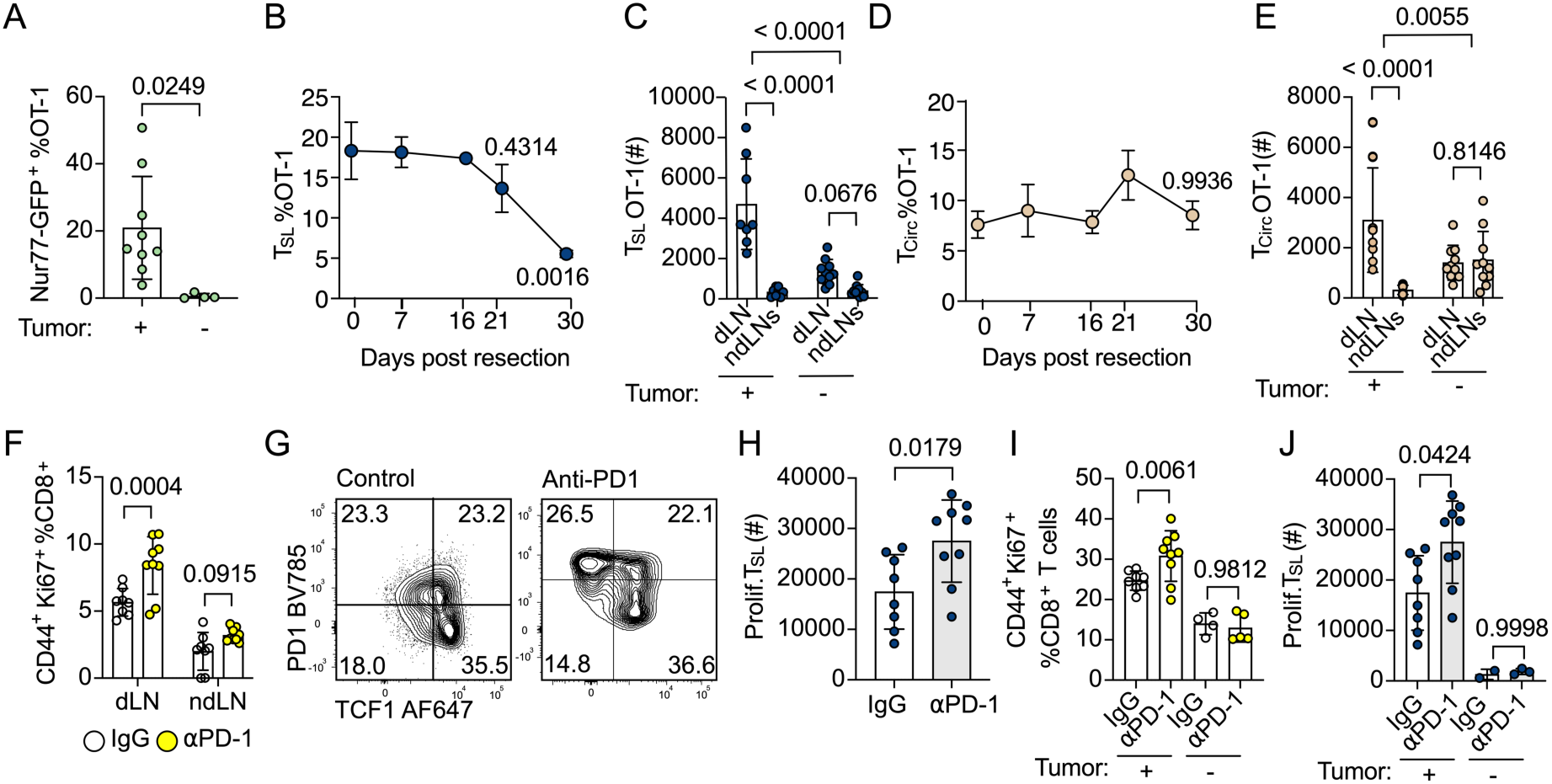
Lymphatic transport maintains the antigen-dependent niche and response to ICB. **(A)** Percent of Nur77-GFP^+^ of OT-1 T cells in the draining lymph node (dLN) of tumor-bearing (+) versus resected (-) mice. YummOVA tumors were analyzed at day 25-28 post-implantation or resected at day 20 and analyzed 16 days later. (**B)** Percent of stem-like (TCF1^+^PD1^INT^, T_SL_) of OT-1 T cells in the dLN over time post-resection. **(C)** Number of T_SL_ OT-1 T cells in dLN and ndLNs at 0- or 30-days post resection. **(D)** Percent of circulating (CD44^+^CD62L^+^, T_Circ_) of OT-1 T cells in dLNs over time post-tumor resection. **(E)** Number of T_Circ_ OT-1 T cells in dLN and non-dLNs (ndLN) at 0- or 30-days post resection. **(F)** Percent of CD44^+^Ki67^+^ of CD8^+^ T cells in the dLN and ndLN in mice treated with IgG or anti-PD1 for 48 hours. **(G)** Representative plot of TCF1 and PD1 expression in CD8^+^ T cells in the dLN of mice treated with IgG or anti-PD1. **(H).** Number of Ki67^+^T_SL_ CD8^+^ T cells dLNs of mice treated with IgG or anti-PD1. **(I)**. Percent of CD44^+^ Ki67^+^ of CD8^+^ T cells in mice treated with IgG or anti-PD1 in tumor (+) and resected (-) dLNs. **(J)**. Number of Ki67^+^T_SL_ in tumor (+) and resected (-) dLNs. For all experiments except (**B, D**), each dot represents a mouse. For **B, D**: (0 n= 8, 7 n=11, 16 n=5, 21 n=10, 30 n=10). Data was analyzed using two-sided, unpaired Student’s t-test **(A, H**) and one-way ANOVA adjusted for multiple comparisons (**B, C, D, E, F, I, J).**

Stem-like cells proliferate following ICB treatment and are required for efficacy in preclinical models^43–49^, with their presence also associated with clinical response in humans^1,48,50^. Accordingly, in our models, PD-1 antibodies triggered a proliferative response in the tumor-draining LNs within 48 hours, with no proliferation seen in non-draining LNs (**Fig. 4F**) or spleen (**Extended Data Fig. 4G**) despite the presence of OT-1 TCR-tg in both locations. PD-1 treatment expands a proliferating stem-like (**Fig. 4G and H**) and induces stem-like T cells to give rise to non-proliferative, effector T cells in the draining LN. The dependence of this early ICB response on stem-like T cells in the tumor-draining LN raises the likelihood that tumor resection, by depleting antigen and the stem-like state, would impair the αPD-1 response. Indeed, tumor resection impaired the 48hr proliferative burst observed upon αPD-1 treatment leading to a reduction in proliferating CD8^+^ T cells (**Fig. 4I**) and a dramatic loss of the proliferating stem-like T cell population (**Fig. 4J**). Importantly, despite their shared TCR and tumor reactivity, memory populations did not proliferate in response to ICB following tumor resection, highlighting the importance of maintaining the LN antigen presentation niche (**Extended Data Fig. 4J and K**) for ICB response.

### Lymph node metastasis disrupts the stem-like T cell niche

Our data indicate that a recirculating effector population repopulates a functional, antigen- and DC-dependent niche in tumor-draining LNs to activate the stem-like state and maintain ICB responsiveness over time. Therefore, the functional status of the lymphatic system, which is often disrupted by standard of care therapy, contributes to the systemic anti-tumor immune potential of the patient. This model has recently been tested with LN sparing regimens in combination with ICB in the clinic^51–55^, leading to better responses compared with ICB after surgery. A major unresolved question, however, is to what extent LN metastasis, which distorts LN architecture, expands regulatory T cells^14,56–60^, and alters DC maturation, positioning, and function^56,57^, compromises the integrity of this dynamic niche and thereby limits ICB efficacy.

To address this question, we first examined metastatic sentinel LNs from a cohort of patients with stage III/IV melanoma using whole-slide, highly multiplexed imaging. Using an automated AI-based nuclear segmentation (HALO), normalization, and clustering pipeline, we systematically classified and manually annotated cellular populations across all samples. In these metastatic LNs, we identified naive (CD8^+^CD45RO^-^TCF1^+^, T_Naive_), exhausted (CD8^+^CD45RO^+^TCF1^−^TOX^+^PD-1^+^, T_EX_), effector (CD8^+^CD45RO^+^GZMB^+^IFNG^+^, T_Eff_), and stem-like T cells (CD8^+^CD45RO^+^TCF1^+^TOX^+^, T_SL_) (**Extended Data Fig. 5A and B**). To analyze if there are differences in the neighborhoods of each T cell state, we identified the 10 nearest neighbors of each T cell and calculated the enrichment (log2 fold change) of neighbor frequencies by cell type as compared to a simulated control^61^. Comparison of the neighborhoods across T cell states revealed that naive T cells were most likely to associate with other naive CD4^+^ and CD8^+^ T cells. Stem-like CD8^+^ T cells were instead proximal to an activated CD4^+^ niche including CD4^+^ICOS^+^PD-1^+^ T cells, reported to be tumor reactive^62,63^, as well as regulatory T cells (CD4^+^FOXP3^+^)^61^. Interestingly, stem-like T cells were also found to be close to mature DCs (CD11C^+^CD40^+^PD-L1^+^MHC-II^+^) (**Fig. 5A and B**), in line with our model that the stem-like state is antigen-dependent, and reminiscent of the CD4:DC:CD8 complexes observed in primary tumors and important for ICB response^64^. Effector CD8^+^ T cells maintained some association with mature DCs but not to CD4^+^ T cells and began to enrich for proximity to CD8^+^ memory populations, immature DCs and antigen-presenting myeloid cells (macrophage and monocytes), while exhausted T cells became further associated with myeloid cells, neutrophils, and tumor (**Fig. 5A and B**). To reveal the largest differences in neighbor frequencies between stem-like and effector T cells and stem-like and exhausted T cells, we performed Cohen’s d Effect size calculations. This confirmed that exhausted/Effector T cells were more likely to be next to MHC-II^+^ myeloid cells while stem-like T cells were more likely to be next to CD4^+^ICOS^+^PD- 1^+^T cells (**Fig. 5C and 5D**). Consistently, we found a stepwise shift away from mature DCs as cells moved from the stem-like state to effector and exhausted, which corresponds with a stepwise increase in association with MHCII^+^ myeloid cells (macrophages and monocytes) (**Fig. 5E and F**) and, as such, an effective switch in proximal antigen-presenting partners. We validated these antigen-presenting/stem-like niches in a second multiplexed imaging data set composed of 21 melanoma LN metastases^65^, again, stem-like T cells were closer to CD4 T cells and DCs than exhausted T cells (**Fig. 5G**). Thus, in metastatic LNs, terminally differentiated cells arise that occupy distinct niches from the regenerative, stem-like population.

**Figure 5.**
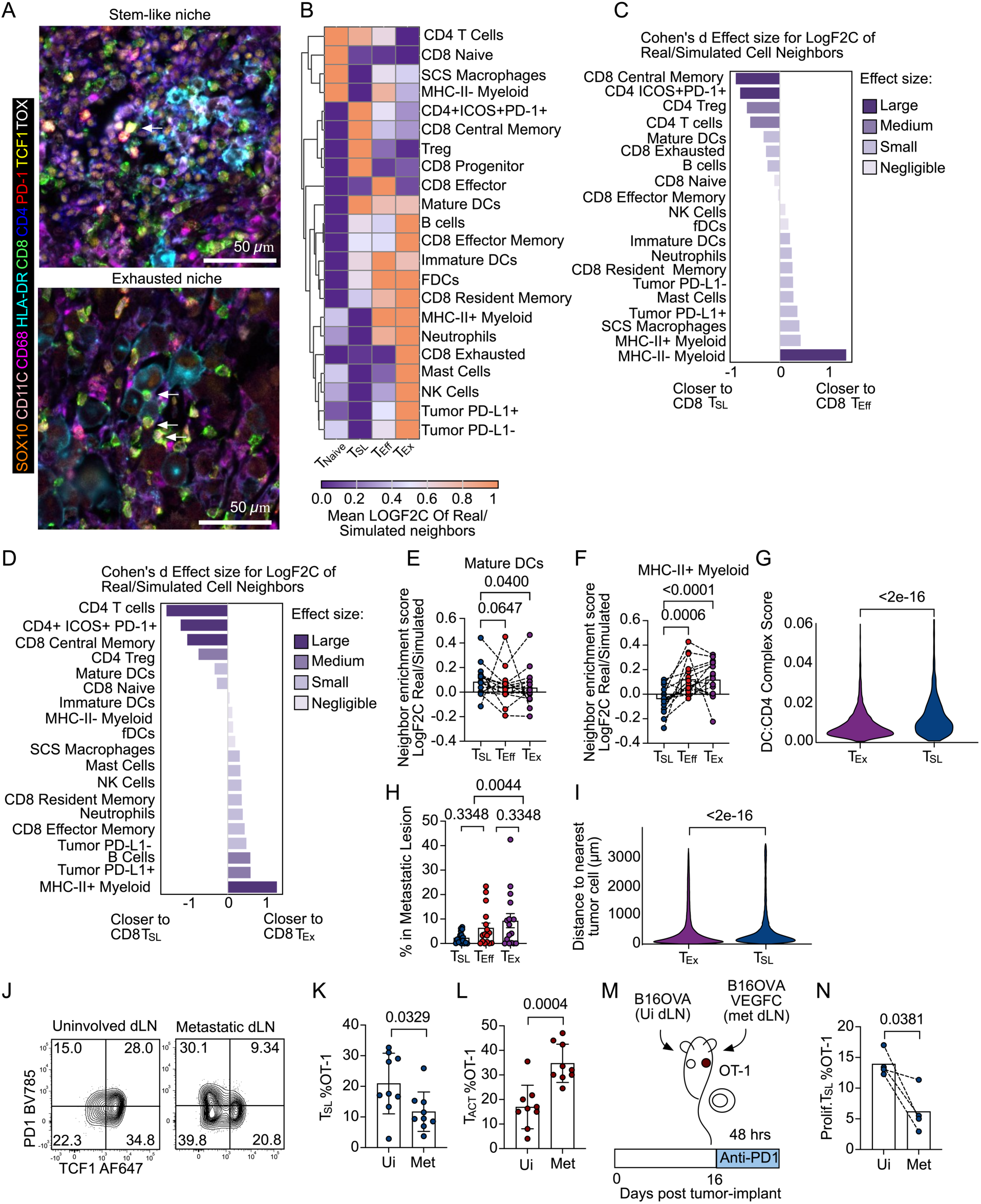
Lymph node metastasis disrupts the stem-like LN niche. **(A)** Representative multiplex fluorescence images of stem-like niche (top) and exhausted niche (bottom) in sentinel lymph nodes (LNs) of human primary melanoma patients. SOX10 melanoma cells (orange), CD11C dendritic cells (light pink), CD68 macrophages (bright pink), MHC-II+ (cyan), CD8 T cells (green), CD4 T cells (blue), PD-1 (red), TCF1 (yellow) and TOX (white) are shown. Arrows indicate presence of stem-like T cells (CD8^+^CD45RO^+^TCF1^+^TOX^+^) (top) and exhausted (CD8^+^CD45RO^+^TCF1^-^TOX^+^PD1^+^) (bottom). Scale bar is 50 μm. **(B)** Heatmap showing mean LOGF2C of Real/Simulated Neighborhoods of Naive (T_Naive_), stem-like (T_SL_), effector (T_Eff_), and exhausted (T_EX_) T cells. Expression normalized by row. **(C)** Cohen’s d effect size for Log2FC of real/simulated cell neighbors comparing T_SL_ to T_Eff_ neighbors. **(D)** Cohen’s d effect size for Log2FC of real/simulated cell neighbors comparing T_SL_ to T_EX_ neighbors. **(E)** Neighbor enrichment score for the specified cell types with mature DCs (CD11C^+^CD40^+^PD-L1^+^MHC-II^+^). **(F)** Neighbor enrichment score for the specified cell types with MHC-II^+^ myeloid cells. (**G**) Complex score (inverse average distance) of the CD8 state with the nearest CD4 (CD4^+^) and DC (CD11C^+^HLA-DR^+^CD14^-^CD68^-^SOX10^-^) (average distance of the three-cell triplet was computed as mean of three pairwise differences). **(H).** Percent of T_SL_, T_Eff_ or T_EX_ in LN metastatic lesion. **(I)** Distance to nearest tumor cell (um) for exhausted (CD8⁺ CD45RO⁻ TCF1⁻ PD1⁺ TOX^+^, T_EX_) or stem-like (CD8⁺ CD45RO⁺ TCF1⁺ TOX⁺, partial PD1⁺, T_SL_) T cells in independent cohort. (**J**). Representative plots of TCF1 and PD1 expression in CD44^+^ OT-1 T cells in draining LNs (dLNs) of B16OVA tumors (uninvolved) and B16OVA VEGFC tumors (metastatic) 18 days post-tumor implantation. (**K**). Percent of stem-like (TCF1^+^PD1^INT^, T_SL_) and (**L**) activated (TCF1^-^PD1^+^, T_ACT_) of OT-1 T cells in uninvolved or metastatic dLNs. **(M)** Schematic of B16OVA/B16OVA VEGFC co-implantation and ICB timeline. **(N)** Percent of KI67^+^ T_SL_ of OT-1 T cells in uninvolved or metastatic dLNs after anti-PD1 treatment. For (**B, E, F**) each dot represents a patient sample (n=16). For (**K, L, N**) each dot represents a mouse. Data was analyzed using Friedman test adjusted for multiple comparisons (**E, F, H**) and using two-sided, unpaired **(K, L**) and paired (**N**) Student’s t-test. For (**G**) data was analyzed using a per-complex Wilcoxon test: compared complex scores of CD8 T_SL_ versus CD8 T_EX_ across all complexes. For (**I**) data was analyzed using a paired Wilcoxon rank-sum test.

Exhausted T cells were also more likely to be proximal to PD-L1^+^ tumors than stem-like T cells (**Fig. 5D**), indicating that the exhausted T cell state was likely driven by the tumor itself, seemingly recapitulating the differentiation trajectory observed in primary tumors. Indeed, while stem-like T cells were predominantly localized outside of the metastatic lesions in these LNs, both effector and exhausted T cells were more likely to be within tumor lesions (**Fig. 5H**). We again validated these findings in the second data set, where exhausted T cells were also closer than stem-like T cells to tumor cells (**Fig. 5I**). These data indicate that the arrival and outgrowth of tumor cells disrupts the antigen presentation landscape leading to local stem-like differentiation into effector and exhausted progeny.

To test this directly, we returned to our preclinical models to ask whether metastasis is sufficient to induce stem-like T cell differentiation in the tumor-draining LN and compromise response to immunotherapy. We predicted that the disruption of antigen presenting partners would force the LN trajectory towards terminal differentiation and thereby deplete the potential for ICB reinvigoration. We compared B16F10 tumors expressing OVA (B16F10.OVA, uninvolved dLN (UI)) to B16F10.OVA tumors also overexpressing the lymphangiogenic growth factor, VEGF-C, which enhances rates of LN metastasis^66^ (B16F10.OVA VEGFC, metastatic dLN (Met)). Analyzing T cell states in the draining LN 18 days post-tumor implantation, a time point sufficient to establish LN metastases^67^, revealed a shift in T cell state (**Fig. 5J**), with a decreased proportion of stem-like (**Fig. 5K**) and the appearance of an activated, effector like OT-I T cell (**Fig. 5L**). These data were aligned with the interpretation that stem-like T cells are induced to terminal differentiation upon metastatic invasion, also seen in our human analyses, due to a shift in their local antigen presentation landscape. To then determine the functional impact of LN metastasis on acute immunotherapy response we treated mice with ICB as done previously (**Fig. 4**). We used the co-implantation experimental set up where both tumors share a systemic anti-OVA immune response but differ in their local microenvironment (uninvolved LN vs. metastatic LN) (**Fig. 5M**), allowing us to test the effects of LN metastasis while controlling for systemic effects. In this setting, despite intact lymphatic transport and increased DC migration from the periphery^66^, the presence of tumor cells within the LN blunted the ICB-induced proliferative stem-like response (**Fig. 5N**). These data collectively argue for the integrity of the lymphatic system, both transport from the periphery and the architectural maintenance of the tumor-draining LN, as a key parameter driving the persistence of a regenerative T cell circuit that fuels the anti-tumor immune response over time.

## Discussion

Successful anti-tumor immunity depends upon both CD8^+^ T cell differentiation into various functional states and the ability to mobilize CD8^+^ T cells to tumors for effective targeting^3–5^. While the contribution of systemic, circulating T cells is evidenced by matching peripheral and tumor clones in patient studies^9^, an understanding of population dynamics over time and specifically how T cell states are compartmentalized anatomically has been challenging to deconvolve from static time points in both mice and humans. Here we leveraged a dynamic, photoconvertible system to track the movement of T cells out of the tumor microenvironment and define a migratory loop that recycles intratumoral effector T cells to sustain an ICB-responsive stem-like pool in tumor-draining LNs and ultimately the effector response in tumors. Constitutive lymphatic transport is required both to recycle tumor-reactive T cells and to chronically load the tumor-draining LN with antigen, together maintaining a dynamic niche that preserves anti-tumor immunity over time.

We demonstrate that effector T cells escape chronic antigen stimulation^24^ and thereby the exhausted differentiation trajectory by egressing from the tumor and undergoing a proliferative burst in the tumor-draining LN. These egressing T cells maintain the stem-like pool that feeds intratumoral effectors, creating a regenerative circuit that sustains anti-tumor immune surveillance. A similar population of chronic effector T cells spans the tumor and LN compartment in patients where its representation in both locations is associated with response to neoadjuvant therapy (Wang et al, co-submitted). We therefore propose that a continuous bi-directional circuit between tumor and LN maintains a more plastic state that confers flexibility in differentiation, while tissue retention locks CD8^+^ T cells into terminal differentiation and dysfunction over time. This population-level plasticity is possibly due in part to intrinsic T cell plasticity. Memory precursors can form from effector cells after antigen clearance through the induction of TCF1, which would help tune memory quantitatively to match the peak of the effector response^36^. Consistently, we see evidence for conversion from an effector state to a memory-like state upon tumor egress. It is also likely that plasticity is maintained at the population level, such that the constitutive movement of less-differentiated populations preserves their functional potential, preventing their sequestration and terminal differentiation at sites of high antigen load. In this way, cells are available to continuously feed into the effector T cell differentiation trajectories observed in mice and patients^19,20^. This model is consistent with observations that KLF2 regulates these effector differentiation pathways^19,20^ and serves as a master regulator of T cell migratory capacity^29–32^, suggesting perhaps that the effects on trafficking are a causal driver of the differentiation potential.

While we find that tumor-egressing effector T cells are sufficient to repopulate LN stem-like T cells, this state transition depends upon antigen reencounter in the LN, indicating that the antigenic status of the LN might direct the differentiation decision. In skin viral infection, where antigen is not presented to T cells after the initial phase of priming (day 5), egressing CD8^+^ T cells are directed towards memory and seed LN resident T cells^23^. In contrast, antigen reencounter in tumor-draining LNs appears to disrupt a memory program, skewing T cells to a stem-like self-renewing state and driving persistence. These observations are aligned with the requirement for chronic antigen to induce and maintain the stem-like state in both tumors^11^ and chronic infection^68,69^, but suggests that the chronicity comes from continuous T cell recirculation and antigen re-encounter through diverse anatomic niches. Indeed, in a recent neoadjuvant ICB melanoma trial, clonally matched T cells in the LN and tumor constituted a chronic effector differentiation trajectory that was associated with response (Wang et al, co-submitted). Functional regional lymphatic transport, which is itself maintained by peripheral tissue cytotoxic responses^67^, ensures the integrity of this circuit by regulating the antigen presentation landscape and limiting metastasis in the LN. Indeed, we find that tumor cell arrival to the LN disrupts the co-localization of T cells with professional DCs and insteads promotes T cell differentiation, which may depend upon engagement with antigen-presenting myeloid and tumor cells. In metastatic LNs, T cell differentiation depletes the stem-like population and thereby inhibits response to ICB. Similar changes in stem-like T cell niche and abundance as a function of metastasis are associated with poor response to ICB in patients^10^.

These data build onto emerging literature that suggest that the integrity of the regional lymphatic basin is key to determining anti-tumor immune efficacy^13,14,59,67^, participating not only in initial priming but importantly in the durable maintenance of anti-tumor immune surveillance over time. This may in part explain why neoadjuvant immunotherapy, where tumor and LN remain intact, yields better responses than adjuvant therapy^51–55^, and may guide clinical management of regional disease in the context of active immunotherapy. Importantly, our data provide a mechanistic framework for understanding recent clinical findings, where response to neoadjuvant αPD1 is associated with a population of migratory, chronic effectors in melanoma (Wang et al, co-submitted), head and neck cancer (Maroun et al, co-submitted), and glioblastoma (Wang et al, co-submitted). Overall, these data suggest that a continuous lymphatic circuit between tumor and tumor-draining LNs recycles and expands less differentiated T cell states associated with treatment response preserving therapeutically relevant clonal differentiation programs over time.

## METHODS

### Mice

C57Bl/6 laboratory mice (*Mus musculus)* (Strain #000664); C57BL/6-Tg(TcraTcrb)1100Mjb/J (OT-1; strain: 003831), and C57BL/6-Tg(Nr4a1 EGFP/cre)820Khog/J (Nur77-GFP; strain: 016617) were purchased directly from Jackson Laboratories. OT-1 Thy1.1 mice were generated by crossing OT-1 mice with B6 Thy1.1 mice (B6.PL-*Thy1^a^*/CyJ, strain: 000406) purchased from Jackson Laboratories. OT1 TCF1 reporter mice were generated by crossing OT1 mice with Tcf7 fl/fl mice (Tcf7^GFP^ flox, Strain #:030909) that were a gift from Kamal Khanna (NYU School of Medicine). B6.Cg-Tg(CAG-tdKaede)-15Utr (Kaede-Tg) were obtained via D.J. Fowell in agreement with RIKEN BioResource Research Center. B6.FVB-1700016L21Rik^Tg(Itgax-HBEGF/EGFP)57Lan^/J (Cd11c-DTR; strain: 004509) were a gift from S.Naik (Mount Sinai). All animal procedures were approved by and performed in accordance with the Institutional Animal Care and Use Committee at NYU.

### Cell culture

The murine melanoma cell lines YummOVA, provided by Ping-Chih Ho (University of Lausanne), and Yummer1.7, provided by M.W. Bosenberg (Yale University), were cultured in DMEM and Ham’s F12 media mixed 1:1 and supplemented with 10% Fetal Bovine Serum (FBS), 1% penicillin/streptomycin (P/S) and 1% Non-essential amino acids. The murine melanoma cell line B16-F10 OVA was provided by M.A. Swartz at University of Chicago and cultured in DMEM media supplemented with 10% FBS and 1% P/S. The B16F10-OVA-VEGFC cells^66^ were passaged in DMEM High Glucose (Cytiva, SH30243.FS) supplemented with 10% FBS and 1% penicillin-streptomycin (Lonza, BW17602E).Cell lines were maintained in incubators at 37C 5% CO_2_. All cell lines are maintained in culture and used for *in vivo* applications at low passages (<25). All lines have been pathogen tested by IDEXX (impact testing) and found to be negative for both mycoplasma and rodent pathogens.

### In vivo tumor growth

One day prior to tumor implant, mice were injected intravenously with 0.4x10^5 (for YummOVA experiments) or 1x10^6 (for B16OVA/B16-OVA-VEGFC experiments) naive splenocytes from OT-1, OT-1 Nur77GFP, OT-1 TCF1^fl/fl^ Ubc-Cre or OT-1 Thy1.1 strain. On the day of injection, mice were shaved and injected with 500K YummOVA, B16OVA or B16OVA VEGFC tumor cells resuspended in isotonic saline intradermally on the upper back. Following injection, mice were monitored daily for health and euthanized on the indicated days.

### Adoptive transfer

Single cell suspensions of OT-1 mice were generated by passing spleens through 70 μM filter cell strainer and lysis of red blood cells was performed with ACK lysis buffer for 1 minute. After CD8 count was determined by flow cytometry staining, calculations were made and splenocytes were transferred i.v. in 200 uLs of PBS.

### In vitro T cell activation

Single cell suspensions of OT-1 mice were generated by passing spleens through 70 μM filter cell strainer and lysis of red blood cells was performed with ACK lysis buffer for 1 minute. Splenocytes were seeded in 12-well plates (6-8x10^6 cells/mL) in RPMI media supplemented with 10% FBS, 1%P/S, 1 nM SIINFEKL and 10 ng/mL of IL-2. After 48 hours, media was replaced with fresh media with SIINFEKL and IL-2. After 72 hours, cells were collected and transferred i.v.

### Diphtheria toxin treatment

On day 20 post tumor-implantation, diphtheria toxin (Sigma-Aldrich, catalog #D0564-1MG) was administered i.p. (4 ng/gr of body weight) every 2 days for 6 days. On day 26, tumors and tumor-draining lymph nodes were collected.

### Kaede-Tg photoconversion

Murine melanoma cell lines were implanted in the back of Kaede-Tg mice. At the indicated time points, mice were photoconverted as previously described^24^. 24 hours after photoconversion, unless otherwise specified, tumors and tumor-draining lymph nodes were collected.

### Survival surgeries

At the indicated time points (10 days or 20 days after tumor implantation), tumors were resected. Surgical garb and sterile gloves, as well as sterile instruments were used. The surgical site was prepared by shaving of hair and surgical betadine scrub alternated with 70% alcohol. Mice were anesthetized during surgery placed in isothermal pad for thermoregulatory support. Recovery from anesthesia took place in a clean cage with a heat source and mice were monitored for signs of distress. Analgesics were provided for 48 hours post-operatively (Carprophen 5mg/kg subcutaneously every 24 hours) and mice were weighed for signs of weight loss.

### In vivo antibody treatment

On day 17, 31 or 25 post-tumor implantation mice were treated with FTY720 (25 grams/mouse) and 200 ug anti-PD1 (RMP1-14, Bio X Cell catalog #BE0146) or IgG control (Sigma-Aldrich, catalog #I8015-100MG). For resected tumors, mice were treated 36- or 47-days post-resection. For low dose anti-CD90.1 experiment, mice were treated on day 13 post tumor implantation with 1 ug of anti Thy1.1 antibody (HIS51, BD, catalog #554892) or IgG (Sigma-Aldrich, catalog #I8015-100MG) control. For treatment with anti-CD62L, mice were treated with 100 ug (Clone: Mel-14, Bio X Cell catalog #BE0021) or IgG (Sigma-Aldrich, catalog #I8015-100MG) at the indicated time points.

### Tissue collection and processing

To create single cell suspensions, LNs were opened with syringes and digested with 1 mg/mL collagenase D and DNase I 80 U/mL for 10 minutes in an incubator at 37C. Suspensions were passed through a 70 μM cell strainer. Spleens were passed through 70 μM cell strainers and red blood cells were lysed with 1 minute treatment with ACK lysis buffer. Tumors were minced and digested with 1 mg/mL collagenase D and DNase I 80 U/mL for 25-30 minutes at 37C with agitation. Suspensions were passed through a 70 μM cell strainer.

### Immunostaining and flow cytometry

Single cell suspensions were first stained with live/dead dye for 15 minutes at 4C in the dark. For tetramer staining, single cell suspensions were then stained with H-2K^b^-OVA_257–264_-BV421 (provided by NIH Tetramer Core) for 45 minutes at room temperature in the dark. For surface staining, cells were incubated with antibodies at the indicated concentrations diluted in PBS 1% BSA for 30 minutes at 4C in the dark. Cells were then washed and fixed with 1% paraformaldehyde solution overnight unless otherwise stated. For intracellular cytokine staining, BD Cytofix/Cytoperm was used following manufacturer instructions. For transcription factor staining, True Nuclear Transcription Buffer set was used following manufacturer instructions. A list of antibodies and dilutions used can be found in Supplementary table 3.

### scRNA-seq analysis

On day 21 after implantation of 500K Yummer1.7 in Kaede-Tg mice (n=3) we photoconverted the tumors and 24 hours later single cell suspensions were enriched for CD45^+^ using a QuadroMACs magnetic separator (CD45-APC, APC beads; Miltenyi Biotec). We stained with DAPI and sorted Kaede-red^+^CD45^+^ cells from the LN (LN) and CD45^+^ cells from the tumor (Tumor) (SY3200 HAPs). Sorted cells were pooled and loaded onto a 10X Genomic Chromium Instrument to generate single-cell gel beads in emulsion. Libraries were prepared using Single Cell 3’ Reagent kits (10x Genomics) and sequenced using Illumina Novaseq 6000.

### Initial data processing

Single-cell RNA-seq data were processed with Cell Ranger (10x Genomics) using the mouse reference genome, and downstream analyses were performed in R using the Seurat package. Cells with fewer than 200 or more than 2,500 detected features, or with >5% of reads mapping to mitochondrial genes, were excluded. LN and tumor samples were merged using the **merge** function. Gene expression values were normalized using the **LogNormalize** method (scale factor = 10,000), and the top 2,000 variable features were identified with **FindVariableFeatures**. The data were then scaled using **ScaleData**. The number of principal components (PCs) to retain was determined based on inspection of elbow plots and JackStraw plots.

### Filtering CD8^+^ T cells

To analyze CD8 T cells, we used Seurat to subset cells expressing T cell receptor–related genes (*Cd3d*, *Cd3e*, *Trac*, and *Trbc1*). Dimensionality reduction and clustering were performed on the previously computed principal components. We then further restricted the dataset to subclusters expressing *Cd8a*. Final dimensionality reduction and clustering were conducted using 13 PCs and a clustering resolution of 0.5, selected via hyperparameter tuning with **FindNeighbors** and **FindClusters**.

### Scoring of cells

Cells were scored for gene signatures using **AddModuleScore**. A list of all signature gene lists can be found in supplementary table 2.

### Statistical analysis for preclinical data

Statistical analyses were performed using GraphPad Prism (version 10.3) and the stats package in R (version 4.3.2) for sequencing data. The specific statistical tests used for each experiment are reported in the corresponding figure legends. Data were assumed to follow a normal distribution; this assumption was not formally tested. No statistical methods were used to predetermine sample sizes; sample sizes were chosen based on previous publications using similar experimental designs. For in vitro experiments, cells from pooled cultures were randomly assigned to experimental groups. For in vivo experiments, age- and sex-matched mice were randomly assigned to experimental groups. For therapeutic intervention studies, treatments were normalized to tumor volume at beginning of treatment. Investigators were blinded to group allocation for all tumor growth measurements. Mice were excluded from tumor studies if tumors failed to establish, became ulcerated during treatment, or if mice developed signs of extreme lethargy, in accordance with pre-established humane endpoints.

### Clinical data analysis

Metastatic sentinel lymph nodes were collected from patients with cutaneous and acral melanoma (n=16, from 12 patients). Validation cohort consisted in the sub setting of LN metastatic samples (n=21, from 12 patients) from an independent NYU cohort^65^. All patients signed written informed consent for the use of their biospecimens for research and participation in this study. This study was approved by the NYU Langone Health Institutional Review Board-approved consent protocol (IRB #10362). Additional information can be found in Supplementary Table 4.

### Fusion multiplex imaging

Whole-slide Quanterix Fusion multiplex imaging was performed^70^ on 7µm FFPE slides from metastatic sentinel lymph nodes (n=16) from patients with cutaneous and acral melanoma (n=12). Slides with multiple sections or patients with multiple metastatic lymph nodes were analyzed separately. Our 45-plex panel includes DAPI and markers for melanoma cells (SOX10, S100b, CSPG4, NGFR, Axl, GP100), stromal populations (aSMA, PDPN, AQP1, CD31, B-catenin, panCK, Collagen IV, PNAd), and immune cells (CD20, CD8, HLA-DR, CD4, ICOS, Mac2/Galectin-3, CD68, CD3e, CD103, FOXP3, CD45RO, TCF-1, CD14, CD57, IDO1, CD21, CD11c, TOX, CD40, CD1c, Clec9a, tryptase, CD169, CD66a/c/e), as well as cytokines and functional markers (IL-33, Ki67, PD-1, PD-L1, Granzyme B, IFNg). A list of antibody clones can be found in Supplementary Table 5.

### Initial data processing and cellular annotation

Initial image analysis was performed using the HALO Image Analysis Platform (Indica Labs). Nuclear segmentation was performed using a custom-trained HALO AI Nuclei Seg V2 network, and tumor and non-tumor compartments were distinguished using a HALO AI DenseNet V2 network trained on manually annotated regions. The borders of each lymph node section were manually annotated, and staining artifacts and folded tissue were excluded from analysis. Intensity-based binary thresholds were set for every marker using HighPlex FL v4.3.2. Briefly, a positive intensity threshold and minimum positive cell area threshold were manually set on a representative area. These thresholds were then calculated as a function of marker range and calculated for and normalized to the remaining samples using min-max scaling. Thresholds were finally manually inspected and adjusted as needed. Cellular classification, area, marker intensity, threshold classification, tumor classifier label, and coordinates were exported for downstream analysis.

Data exported from HALO were imported by sample to Python v3.9.21 as AnnData objects. The smallest 1% of cells by cellular area and by nuclear staining intensity were removed and marker intensities were Z-scored within each section. After normalization, the top 1% of cells by number of markers expressed or total marker expression were removed. Objects were then merged into a single AnnData object. Cells were annotated by serial Leiden clustering using lineage markers. Cellular clusters that were either uniformly high or uniformly low for all markers were removed. Detailed cellular subtypes were subsequently manually identified using HALO-calculated thresholds and tumor classifier status.

### Neighborhood analysis

For each cell, the 10 nearest neighbors by centroid-centroid distance were calculated using the **neighborhood_analysis()** function from SPACEc (https://github.com/yuqiyuqitan/SPACEc)^71^ adapted to export the cell-level neighbor matrix in addition to the clustered AnnData object. This matrix was then imported into a custom simulation function written with the aid of NYU-proprietary generative AI that creates a simulated neighbor matrix per sample based on the distribution of cell identities within that sample. This simulation was iterated until the predicted mean nearest neighbors to each cell type changed by less than 1% with additional iterations, and the mean nearest neighbors per cell over each iteration (n=1000) was used as the final simulated control. The average real and simulated nearest neighbors to each cell type was calculated within each sample. The neighbor enrichment score is defined as the log2 fold change of mean real nearest neighbors over mean simulated nearest neighbors. The Cohen’s d effect size is calculated between groups using the mean and pooled standard deviation of these sample level cellular enrichment scores. This analysis is conceptually based on the ESI-map analysis in^61^.

### Distance analysis

#### Distance to tumor cells

To evaluate the spatial relationship between CD8 T cell subtypes and tumor cells in LN metastases, we classified CD8 T cells into stem-like (CD8⁺ CD45RO⁺ TCF1⁺ TOX⁺, partial PD1⁺) and exhausted (CD8⁺ CD45RO⁻ TCF1⁻ PD1⁺ TOX^+^) populations using binary marker expression. Distances from each CD8 T cell to the tumor cell were calculated based on cell centroid coordinates. Distances of all CD8 T cells to their nearest tumor cells were compared between subtypes using a paired Wilcoxon rank-sum test at the sample level, visualized as violin plots.

#### Spatial proximity analysis of CD8 T cell/CD4 T cell/DC complexes

For each CD8 T cell, we identified the nearest CD4 T cell and nearest DC using k-nearest neighbors (FNN::get.knnx). The average distance of the three-cell triplet (CD8–CD4, CD8–DC, CD4–DC) was computed as the mean of the three pairwise distances. This per-CD8 cell metric represents the spatial compactness of the complex. To quantify complex formation, we transformed the average distance between components into a composite score by taking its inverse, with a constant of 1 added to avoid division by zero. In this framework, higher complex scores correspond to tighter and more spatially compact complexes. Data was analyzed using per-complex Wilcoxon test: compared complex scores of CD8_Stem-like vs CD8_Exhausted across all complexes.

## Supporting information

External Data Figures

## Data and code availability

Single-cell RNA sequencing data from draining lymph node photoconverted cells have been deposited in the Gene Expression Omnibus (GEO) under accession number GSE322572 and will be made publicly available upon acceptance. Single-cell RNA sequencing data from tumor-infiltrating lymphocytes are available under accession number GSE218455^24^. Imaging data are available upon request.

## Acknowledgements

The authors acknowledge technical assistance and feedback from Brian Quartey and the rest of the Lund lab. This work was supported by grants from the Cancer Research Institute (Lloyd J. Old STAR Award to AWL), the American Association for Cancer Research (AACR-BMS Midcareer Female Investigator Award, AWL), and the NIH (R01CA238163, A.W.L.; R01CA304163A, A.W.L; U54CA263001, AWL, IO; P50CA016087 to AWL and IO). KSV was supported by the NIH (F30CA288142-01A), TM from the Cancer Research Institute (CRI Dr. Keith Landesman Memorial Postdoctoral Fellow (CRI14497), and TAH from the NIH (T32-AI100853).

## Author contributions

Conceptualization: ID, AWL

Methodology: ID, KSV, TM, MI, TAH, MMS

Investigation: ID, KSV, NRB, TM, MI, GW, TAH, MMS

Funding acquisition: AWL

Reagents and Biospecimens: ACH, IO

Supervision: ACH, MS, AWL

Writing – original draft: ID, AWL

Writing – review & editing: ID, KSV, NRB, TM, GW, MI, TAH, MMS, ACH, MS, IO, AWL

## Declaration of generative AI and AI-assisted technologies in the manuscript preparation process

During the preparation of this work the authors used NYU-proprietary generative AI to check grammar, clarity and consistency of text. After using this tool/service, the authors reviewed and edited the content as needed and take full responsibility for the content of the published article.

## References

1. Jansen, C. S. et al. An intra-tumoral niche maintains and differentiates stem-like CD8 T cells. Nature 576, 465–470 (2019).

2. Im, S. J. et al. Characteristics and anatomic location of PD-1+TCF1+ stem-like CD8 T cells in chronic viral infection and cancer. Proceedings of the National Academy of Sciences 120, e2221985120 (2023).

3. Wijesinghe, S. K. M. et al. Lymph-node-derived stem-like but not tumor-tissue-resident CD8+ T cells fuel anticancer immunity. Nat Immunol 26, 1367–1383 (2025).

4. Prokhnevska, N. et al. CD8+ T cell activation in cancer comprises an initial activation phase in lymph nodes followed by effector differentiation within the tumor. Immunity 56, 107–124.e5 (2023).

5. Connolly, K. A. et al. A reservoir of stem-like CD8+ T cells in the tumor-draining lymph node preserves the ongoing antitumor immune response. Sci Immunol 6, eabg7836 (2021).

6. Huang, Q. et al. The primordial differentiation of tumor-specific memory CD8+ T cells as bona fide responders to PD-1/PD-L1 blockade in draining lymph nodes. Cell 185, 4049–4066.e25 (2022).

7. Li, G. et al. TGF-β-dependent lymphoid tissue residency of stem-like T cells limits response to tumor vaccine. Nat Commun 13, 6043 (2022).

8. Schenkel, J. M. et al. Conventional type I dendritic cells maintain a reservoir of proliferative tumor-antigen specific TCF-1+ CD8+ T cells in tumor-draining lymph nodes. Immunity 54, 2338–2353.e6 (2021).

9. Pai, J. A. et al. Lineage tracing reveals clonal progenitors and long-term persistence of tumor-specific T cells during immune checkpoint blockade. Cancer Cell 41, 776–790.e7 (2023).

10. Rahim, M. K. et al. Dynamic CD8+ T cell responses to cancer immunotherapy in human regional lymph nodes are disrupted in metastatic lymph nodes. Cell 186, 1127–1143.e18 (2023).

11. Hor, J. L. et al. Inhibitory PD-1 axis maintains high-avidity stem-like CD8+ T cells. Nature 649, 194–204 (2026).

12. Dähling, S. et al. Type 1 conventional dendritic cells maintain and guide the differentiation of precursors of exhausted T cells in distinct cellular niches. Immunity 55, 656–670.e8 (2022).

13. Delclaux, I., Ventre, K. S., Jones, D. & Lund, A. W. The tumor-draining lymph node as a reservoir for systemic immune surveillance. Trends Cancer (2023).

14. Reticker-Flynn, N. E. et al. Lymph node colonization induces tumor-immune tolerance to promote distant metastasis. Cell 185, 1924–1942.e23 (2022).

15. Bai, A., Hu, H., Yeung, M. & Chen, J. Krüppel-Like Factor 2 Controls T Cell Trafficking by Activating L-Selectin (CD62L) and Sphingosine-1-Phosphate Receptor 1 Transcription1. J Immunol 178, 7632–7639 (2007).

16. Skon, C. N. et al. Transcriptional downregulation of S1pr1 is required for the establishment of resident memory CD8+ T cells. Nat Immunol 14, 1285–1293 (2013).

17. Takada, K. et al. Kruppel-Like Factor 2 Is Required for Trafficking but Not Quiescence in Postactivated T Cells. J Immunol 186, 775–783 (2011).

18. Carlson, C. M. et al. Kruppel-like factor 2 regulates thymocyte and T-cell migration. Nature 442, 299–302 (2006).

19. Fagerberg, E. et al. KLF2 maintains lineage fidelity and suppresses CD8 T cell exhaustion during acute LCMV infection. Science 387, eadn2337 (2025).

20. Tsui, C. et al. Lymph nodes fuel KLF2-dependent effector CD8+ T cell differentiation during chronic infection and checkpoint blockade. Nat Immunol 26, 1752–1765 (2025).

21. Shen, J. et al. Krüppel-like factors 2 and 3 regulate T cell exhaustion by directing T cell residency and migration. Immunity S1074761326000919 (2026) doi:10.1016/j.immuni.2026.02.021.

22. Beura, L. K. et al. T Cells in Nonlymphoid Tissues Give Rise to Lymph-Node-Resident Memory T Cells. Immunity 48, 327–338.e5 (2018).

23. Heim, T. A. et al. Lymphatic vessel transit seeds cytotoxic resident memory T cells in skin draining lymph nodes. Science Immunology 9, eadk8141 (2024).

24. Steele, M. M. et al. T cell egress via lymphatic vessels is tuned by antigen encounter and limits tumor control. Nat Immunol 24, 664–675 (2023).

25. Li, Z. et al. In vivo labeling reveals continuous trafficking of TCF-1+ T cells between tumor and lymphoid tissue. J Exp Med 219, e20210749 (2022).

26. Debes, G. F. et al. Chemokine receptor CCR7 required for T lymphocyte exit from peripheral tissues. Nat. Immunol. 6, 889–894 (2005).

27. Bromley, S. K., Thomas, S. Y. & Luster, A. D. Chemokine receptor CCR7 guides T cell exit from peripheral tissues and entry into afferent lymphatics. Nature Immunology 6, 895–901 (2005).

28. Galván-Peña, S., Zhu, Y., Hanna, B. S., Mathis, D. & Benoist, C. A dynamic atlas of immunocyte migration from the gut. Sci. Immunol. 9, eadi0672 (2024).

29. Molodtsov, A. K. et al. Resident memory CD8+ T cells in regional lymph nodes mediate immunity to metastatic melanoma. Immunity 54, 2117–2132.e7 (2021).

30. Takahashi, M. et al. Intratumoral antigen signaling traps CD8+ T cells to confine exhaustion to the tumor site. Science Immunology 9, eade2094 (2024).

31. Li, Z. et al. In vivo labeling reveals continuous trafficking of TCF-1+ T cells between tumor and lymphoid tissue. Journal of Experimental Medicine 219, e20210749 (2022).

32. Macatonia, S. E., Knight, S. C., Edwards, A. J., Griffiths, S. & Fryer, P. Localization of antigen on lymph node dendritic cells after exposure to the contact sensitizer fluorescein isothiocyanate. Functional and morphological studies. The Journal of experimental medicine 166, 1654–1667 (1987).

33. Luckey, C. J. et al. Memory T and memory B cells share a transcriptional program of self-renewal with long-term hematopoietic stem cells. Proceedings of the National Academy of Sciences 103, 3304–3309 (2006).

34. Kaech, S. M., Hemby, S., Kersh, E. & Ahmed, R. Molecular and Functional Profiling of Memory CD8 T Cell Differentiation. Cell 111, 837–851 (2002).

35. Joshi, N. S. et al. Inflammation Directs Memory Precursor and Short-Lived Effector CD8+ T Cell Fates via the Graded Expression of T-bet Transcription Factor. Immunity 27, 281–295 (2007).

36. Abadie, K. et al. Reversible, tunable epigenetic silencing of TCF1 generates flexibility in the T cell memory decision. Immunity 57, 271–286.e13 (2024).

37. Mempel, T. R., Henrickson, S. E. & von Andrian, U. H. T-cell priming by dendritic cells in lymph nodes occurs in three distinct phases. Nature 427, 154–159 (2004).

38. Khan, T. N., Mooster, J. L., Kilgore, A. M., Osborn, J. F. & Nolz, J. C. Local antigen in nonlymphoid tissue promotes resident memory CD8+ T cell formation during viral infection. Journal of Experimental Medicine 213, 951–966 (2016).

39. Wang, G. et al. Antigen-specific profiling identifies T-bet+ melanoma-specific CD8+ T cells associated with response to neoadjuvant PD-1 blockade. Cancer Cell 44, 221–234.e5 (2026).

40. Loo, C. P. et al. Lymphatic Vessels Balance Viral Dissemination and Immune Activation following Cutaneous Viral Infection. Cell Reports 20, 3176–3187 (2017).

41. Roberts, E. W. et al. Critical Role for CD103+/CD141+ Dendritic Cells Bearing CCR7 for Tumor Antigen Trafficking and Priming of T Cell Immunity in Melanoma. Cancer Cell 30, 324–336 (2016).

42. Lund, A. W. et al. Lymphatic vessels regulate immune microenvironments in human and murine melanoma. J Clin Invest 126, 3389–3402 (2016).

43. Huang, A. C. et al. A single dose of neoadjuvant PD-1 blockade predicts clinical outcomes in resectable melanoma. Nat Med 25, 454–461 (2019).

44. Im, S. J. et al. Defining CD8+ T cells that provide the proliferative burst after PD-1 therapy. Nature 537, 417–421 (2016).

45. Utzschneider, D. T. et al. T Cell Factor 1-Expressing Memory-like CD8+ T Cells Sustain the Immune Response to Chronic Viral Infections. Immunity 45, 415–427 (2016).

46. He, R. et al. Follicular CXCR5-expressing CD8+ T cells curtail chronic viral infection. Nature 537, 412–416 (2016).

47. Miller, B. C. et al. Subsets of exhausted CD8+ T cells differentially mediate tumor control and respond to checkpoint blockade. Nat Immunol 20, 326–336 (2019).

48. Siddiqui, I. et al. Intratumoral Tcf1+PD-1+CD8+ T Cells with Stem-like Properties Promote Tumor Control in Response to Vaccination and Checkpoint Blockade Immunotherapy. Immunity 50, 195–211.e10 (2019).

49. Kurtulus, S. et al. Checkpoint Blockade Immunotherapy Induces Dynamic Changes in PD-1−CD8+ Tumor-Infiltrating T Cells. Immunity 50, 181–194.e6 (2019).

50. Sade-Feldman, M. et al. Defining T Cell States Associated with Response to Checkpoint Immunotherapy in Melanoma. Cell 175, 998–1013.e20 (2018).

51. Patel, S. P. et al. Neoadjuvant–Adjuvant or Adjuvant-Only Pembrolizumab in Advanced Melanoma. New England Journal of Medicine 388, 813–823 (2023).

52. Blank, C. U. et al. Neoadjuvant Nivolumab and Ipilimumab in Resectable Stage III Melanoma. New England Journal of Medicine 391, 1696–1708 (2024).

53. Wakelee, H. et al. Perioperative Pembrolizumab for Early-Stage Non–Small-Cell Lung Cancer. New England Journal of Medicine 389, 491–503 (2023).

54. Forde, P. M. et al. Neoadjuvant Nivolumab plus Chemotherapy in Resectable Lung Cancer. N Engl J Med 386, 1973–1985 (2022).

55. Provencio, M. et al. Perioperative Nivolumab and Chemotherapy in Stage III Non–Small-Cell Lung Cancer. New England Journal of Medicine 389, 504–513 (2023).

56. van den Hout, M. F. C. M., et al. Melanoma Sequentially Suppresses Different DC Subsets in the Sentinel Lymph Node, Affecting Disease Spread and Recurrence. Cancer Immunol Res 5, 969–977 (2017).

57. van Pul, K. M. et al. Selectively hampered activation of lymph node-resident dendritic cells precedes profound T cell suppression and metastatic spread in the breast cancer sentinel lymph node. J Immunother Cancer 7, 133 (2019).

58. Kos, K. et al. Tumor-educated Tregs drive organ-specific metastasis in breast cancer by impairing NK cells in the lymph node niche. Cell Rep 38, 110447 (2022).

59. Lei, P.-J. et al. Cancer cell plasticity and MHC-II–mediated immune tolerance promote breast cancer metastasis to lymph nodes. Journal of Experimental Medicine 220, e20221847 (2023).

60. Núñez, N. G. et al. Tumor invasion in draining lymph nodes is associated with Treg accumulation in breast cancer patients. Nat Commun 11, 3272 (2020).

61. Solis, S. M. et al. Cellular interactions in the sentinel lymph node predict melanoma recurrence. 2025.12.15.694104 Preprint at 10.64898/2025.12.15.694104 (2025).

62. Duhen, R. et al. PD-1 and ICOS coexpression identifies tumor-reactive CD4^+^ T cells in human solid tumors. J Clin Invest 132, (2022).

63. Alspach, E. et al. MHC-II neoantigens shape tumour immunity and response to immunotherapy. Nature 574, 696–701 (2019).

64. Espinosa-Carrasco, G. et al. Intratumoral immune triads are required for immunotherapy-mediated elimination of solid tumors. Cancer Cell 42, 1202–1216.e8 (2024).

65. Ibrahim, M. et al. NF1 Loss Remodels Tumor Niches for Immune Evasion. 2026.01.11.698818 Preprint at 10.64898/2026.01.11.698818 (2026).

66. Lund, A. W. et al. VEGF-C promotes immune tolerance in B16 melanomas and cross-presentation of tumor antigen by lymph node lymphatics. Cell Rep 1, 191–199 (2012).

67. Karakousi, T. et al. IFNγ-dependent metabolic reprogramming restrains an immature, pro-metastatic lymphatic state in melanoma. Cancer Cell 10.1016/j.ccell.2025.12.019 (2026) doi:10.1016/j.ccell.2025.12.019.

68. McManus, D. T. et al. An early precursor CD8+ T cell that adapts to acute or chronic viral infection. Nature 640, 772–781 (2025).

69. Chu, T. et al. Precursors of exhausted T cells are pre-emptively formed in acute infection. Nature 640, 782–792 (2025).

70. Goltsev, Y. et al. Deep Profiling of Mouse Splenic Architecture with CODEX Multiplexed Imaging. Cell 174, 968–981.e15 (2018).

71. Tan, Y. et al. SPACEc: a streamlined, interactive Python workflow for multiplexed image processing and analysis. Nat Commun 16, 10652 (2025).

