## Supplementary material for "Lymphatic egress recycles tumor-experienced effector CD8 T cells to sustain immune surveillance": External Data Figures

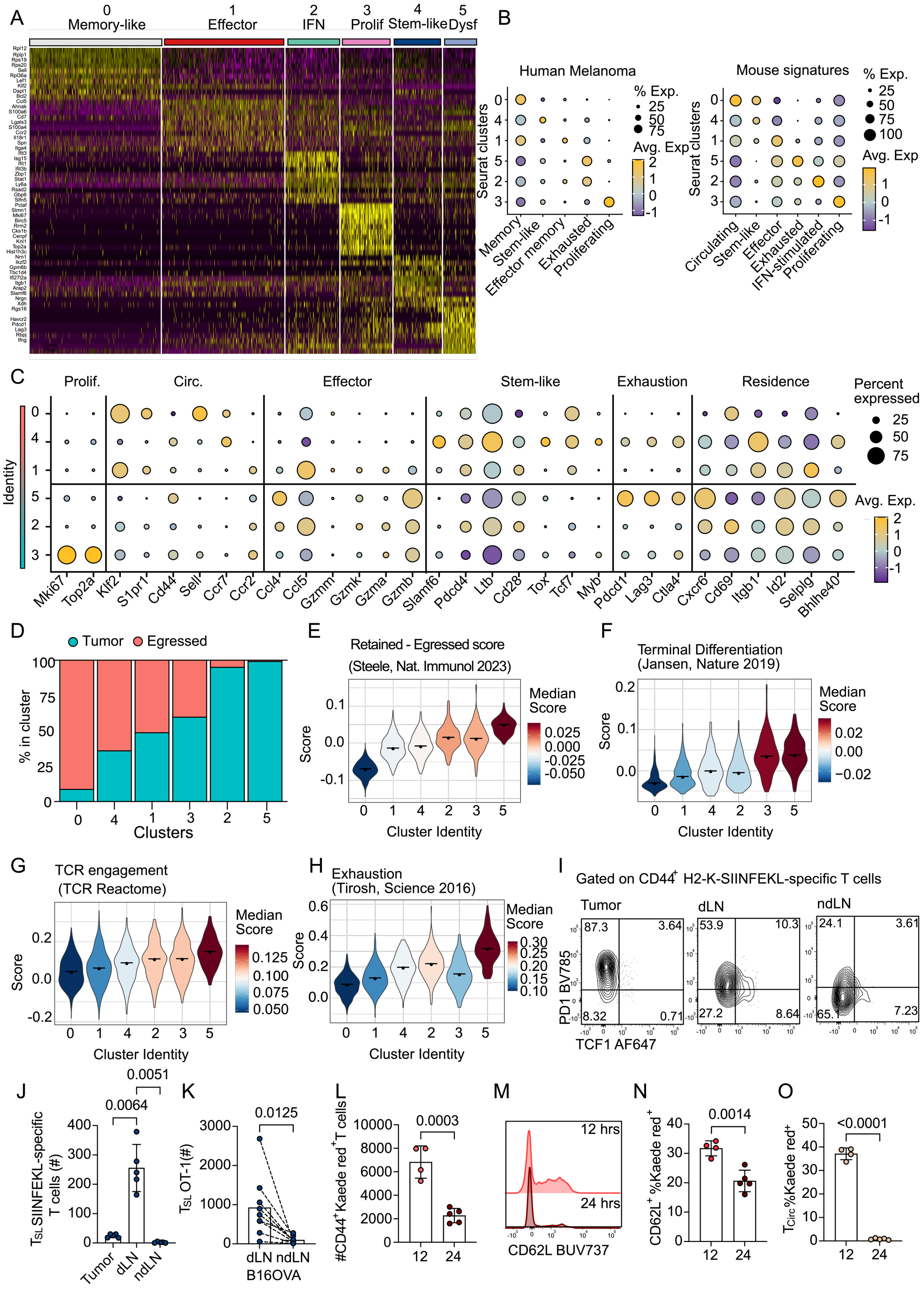


**External Data Figure 1 (Related to Figure 1). Tumor-specific early effector T cells egress tumor microenvironments (A)** Top 10 cluster-defining genes from scRNAseq data. **(B)** Expression of circulating[1], stem-like, effector, exhausted[2] IFN-stimulated and proliferating[3] module scores from mice (left) and memory, stem-like, effector memory, exhausted, proliferating module scores from human melanoma[4] (right) **(C)** Expression of select genes across clusters. **(D)** Proportion of tumor and egressed (Kaede red$^+$ cells in the draining lymph node - dLN) CD8$^+$ T cells by clusters. **(E-H)** Clusters scored by a retained versus egress (expression of retained program–egress program) **(E)**[5], a terminal differentiation[6], **(F)** a TCR engagement (Reactome, R-MMU-202424) **(G)** and an exhaustion[7] signature **(H)** **(I)** Representative plot and **(J)** quantification of TCF1 and PD1 in CD44$^+$ H2-K$^b$-SIINFEKL-specific T cells 26 days post YummOVA implant in tumor, dLN and ndLN and. **(K)** Number of stem-like T cells (TCF1$^+$PD1$^{INT}$, $T_{SL}$) in B16OVA dLN and non-dLN (ndLN) 18 days post tumor implant **(L)** Number of CD44$^+$Kaede red$^+$ CD8$^+$ T cells in dLN 12 and 24 hours post-photoconversion of YUMMER1.7 tumors. **(M)** Representative histogram and **(N)** percent of CD62L expression in Kaede red$^+$ CD44$^+$CD8$^+$ T cells in the dLN 12 and 24 hours after photoconversion. **(O)** Percent of circulating (CD62L$^+$, $T_{CIRC}$) of Kaede red$^+$ cells in the dLN 12 and 24 hours post-photoconversion. (**I-O**) each dot is a mouse. Data was analyzed using two-sided, paired (**K**) and unpaired (**L, N, O**) Student's t-test and one-way ANOVA adjusted for multiple comparisons (**J**)

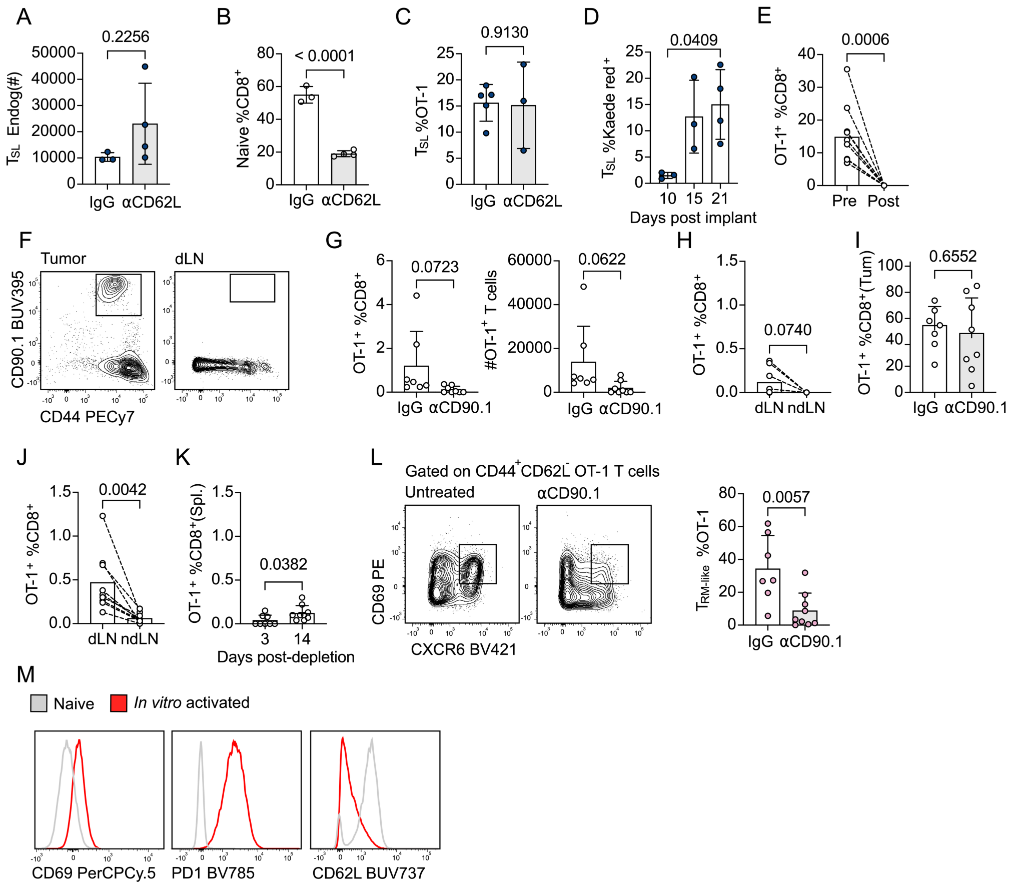


**External Data Figure 2 (Related to Figure 2). Study of T cell trafficking dynamics by blocking entry into lymph node from circulation or depletion of circulating T cells. (A)** Number of endogenous stem-like T cells (CD44$^+$ TCF1$^+$ PD1$^{INT}$, $T_{SL}$) T cells in YummOVA draining lymph nodes (dLNs) treated with IgG or αCD62L. **(B)** Percent of naïve CD8$^+$ T cells in dLNs treated with IgG or αCD62L. **(C)** Percent of $T_{SL}$ of CD44$^+$ OT-1 T cells in B16OVA dLNs treated with IgG or αCD62L. **(D)** Percent of $T_{SL}$ of CD44$^+$ Kaede red$^+$CD8$^+$ T cells in dLNs 24 hours after photoconversion at the indicated time points post-YUMMER1.7 implantation. **(E)** Percent of OT-1 CD8$^+$ T cells in the blood pre and post αCD90.1 treatment. **(F)** Representative plots showing gating of CD90.1 OT-1 T cells in tumor and dLNs 3 days post αCD90.1 treatment. **(G)** Percent and numbers of OT-1 T cells in dLNs 3 days after IgG or αCD90.1 treatment. **(H)** Percent of OT-1 T cells in dLN and non-dLN (ndLN) 3 days post αCD90.1 treatment. **(I)** Percent of OT-1 T cells in tumors (Tum.) 3 days post αCD90.1 treatment. **(J)** Percent of OT-1 T cells in dLN and ndLN 14 days post αCD90.1 treatment. **(K)** Percent of OT-1 T cells in spleens (Spl.) 3- and 14-days post αCD90.1 treatment. **(L)** Representative plots (left) and percent (right) of CD69 and CXCR6 staining in OT-1 CD44$^+$CD62L$^-$ OT-1 T cells 14 days post IgG or αCD90.1 treatment. **(M)** Representative histograms of indicated markers in naïve OT-1 (grey) or *in vitro* activated OT-1 T cells (red) For all data, each dot represents a mouse. Data was analyzed using two-sided, unpaired (**A, B, C, G, I, K, L**) and paired (**E, H, J**) Student's t-test and one-way ANOVA adjusted for multiple comparisons (**D**)

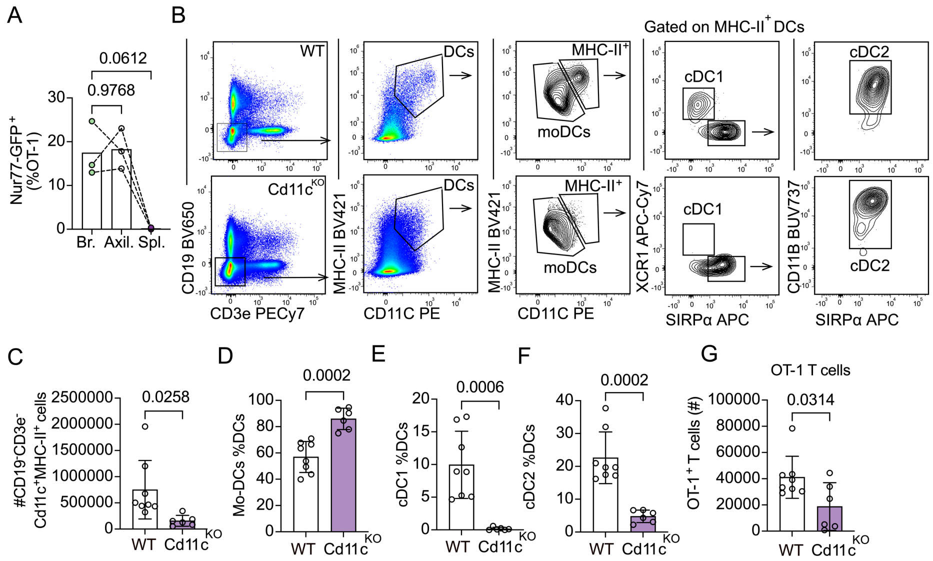


**External Data Figure 3 (Related to Figure 3). Dendritic cell subsets upon diphtheria depletion in tumor models. (A)** Percent of Nur-77GFP$^+$ expression of OT-1 T cells in brachial ipsilateral (Br.), axillary ipsilateral (Ax.) draining lymph nodes (dLN) and spleen (Spl.) **(B)** Representative gating scheme for monocyte-derived DCs (MHC-II$^{LOW}$, moDCs), cDC1s (MHC-II$^+$ Xcr1$^+$) and cDC2s (MHC-II$^+$ Sirp$\alpha^+$ Cd11b$^+$) in Cd11c$^{WT}$ (WT) or Cd11c$^{DTR}$ (Cd11c$^{KO}$) mice treated with diphtheria toxin (DT) **(C)** Number of CD19$^-$CD3$^-$Cd11c$^+$MHC-II$^+$ cells in WT or Cd11c$^{KO}$ **(D-F)** Percent of mo-DCs **(D),** cDC1s **(E)** and cDC2s **(F)** in WT or Cd11c$^{KO}$ mice. **(G)** Number of OT-1 T cells in WT or Cd11c$^{KO}$ mice. For all data each dot represents a mouse. Data was analyzed using two-sided, paired (**A**) or unpaired (**C-G**) Student's t-test.

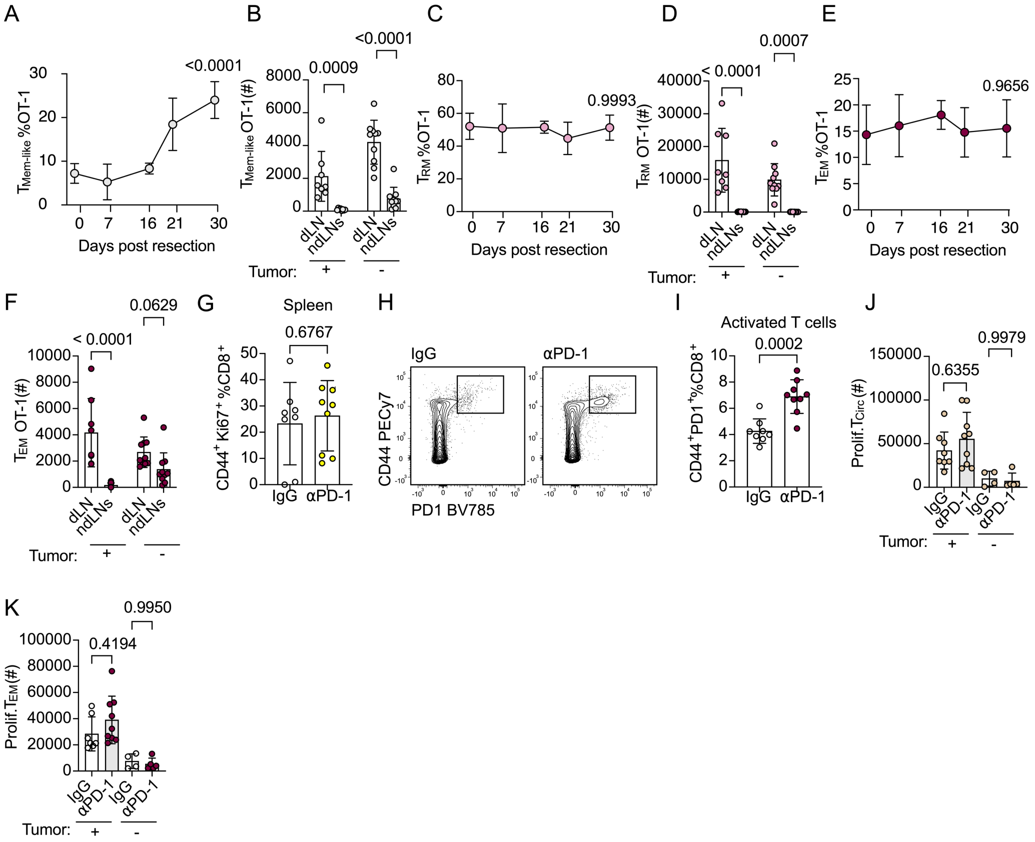


**External Data Figure 4 (Related to Figure 4). Memory T cells equilibrate post tumor resection but do not respond to ICB. (A)** Percent of memory-like ($CD62L^{-}TCF1^{+}PD1^{-}$, $T_{Mem\text{-}like}$) of OT-1 T cells in the draining lymph node (dLN) at the indicated time-points post-tumor resection. **(B)** Number of $T_{Mem\text{-}like}$ OT-1 T cells in dLN and non-dLN (ndLN) at 0- or 30-days post resection**. (C)** Percent of tissue resident memory ($CD62L^{-}CD69^{+}CXCR6^{+}$ $TCF1^{-}$ , $T_{RM}$) of OT-1 T cells in the dLN at the indicated time-points post-tumor resection. **(D)** Number of $T_{RM}$ OT-1 T cells in dLN and ndLNs at at 0- or 30-days post resection**. (E)** Percent of effector memory ($CD62L^{-}CD69^{-}$, $T_{EM}$) of OT-1 T cells in the dLN at the indicated time-points post-tumor resection. **(F)** Number of $T_{EM}$ OT-1 T cells in dLN and ndLNs at at 0- or 30-days post resection**. (G)** Percentage of $CD44^{+}Ki67^{+}$ of $CD8^{+}$ T cells in spleen in IgG or anti-PD1 treated mice. **(H)** Representative plots and **(I)** percent of $PD1^{+}CD44^{+}$ of $CD8^{+}$ T cells in IgG or anti-PD1 treated mice. **(J)** Number of proliferating circulating ($CD44^{+}CD62L^{+}Ki67^{+}$, Prolif. $T_{CIRC}$) and (**K**) $Ki67^{+}$ $T_{EM}$ T cells in dLNs of IgG or anti-PD1 treated mice at day 20 post tumor implant (+) or 36/47 days post resection (-) For all experiments except (**A, C, D**), each dot represents a mouse. For **A, C, D**: (0 n= 8, 7 n=11, 16 n=5, 21 n=10, 30 n=10) Data was analyzed using one-way ANOVA adjusted for multiple comparisons (**A-F, J, K**) and unpaired Student's t-test (**G, I**)

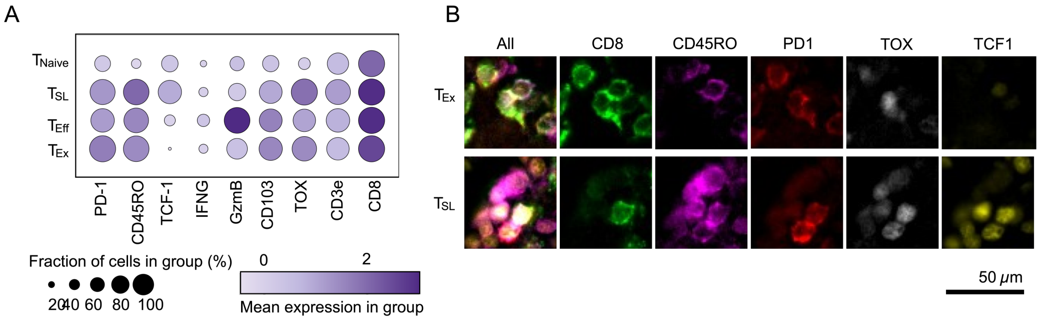


**External Data Figure 5 (Related to Figure 5). Identification of T cell subsets in multiplexed images. (A)** Protein expression of select markers in segmented T cell populations from multiplexed fluorescence imaging. **(B)** Representative multiplex fluorescence images of defining T cell markers: CD8 T cells (green), CD45RO (pink), PD-1 (red), TOX (white) and TCF1 (yellow) Scale bar is 50 μm.

### References


1. Heim, T. A. *et al.* Lymphatic vessel transit seeds cytotoxic resident memory T cells in skin draining lymph nodes. *Science Immunology* **9**, eadk8141 (2024)
2. Miller, B. C. *et al.* Subsets of exhausted CD8+ T cells differentially mediate tumor control and respond to checkpoint blockade. *Nat Immunol* **20**, 326–336 (2019)
3. Giles, J. R. *et al.* Shared and distinct biological circuits in effector, memory and exhausted CD8+ T cells revealed by temporal single-cell transcriptomics and epigenetics. *Nat Immunol* **23**, 1600–1613 (2022)
4. Oliveira, G. *et al.* Phenotype, specificity and avidity of antitumour CD8+ T cells in melanoma. *Nature* **596**, 119–125 (2021)
5. Steele, M. M. *et al.* T cell egress via lymphatic vessels is tuned by antigen encounter and limits tumor control. *Nat Immunol* **24**, 664–675 (2023)
6. Jansen, C. S. *et al.* An intra-tumoral niche maintains and differentiates stem-like CD8 T cells. *Nature* **576**, 465–470 (2019)
7. Dissecting the multicellular ecosystem of metastatic melanoma by single-cell RNA-seq | Science. https://www.science.org/doi/10.1126/science.aad0501.
