## Supplemental Table 4 for "Lymphatic egress recycles tumor-experienced effector CD8 T cells to sustain immune surveillance"

### Table S4. Patient cohort information

#### Metastatic sentinel lymph nodes from cutaneous and acral melanoma patients.

| Primary melanoma anatomic location | Gender | Age at Excision | Stage at Excision | Type | Primary tumor thickness | Mutation status | Treatment before excision |
|---|---|---|---|---|---|---|---|
| Right Upper Back | Female | 62 | Stage IIIC | Cutaneous | 6 | BRAF V600E, ALK T680I | No |
| Left upper back | Male | 69 | Stage IIIC | Cutaneous | 3.1 | BRAF V600K, FGFR1 I234T, FGFR2 S372F | No |
| Left lower back | Male | 62 | Stage IIIC | Cutaneous | 4.1 | BRAF V600E, ALK T680I | No |
| Left upper back | Male | 81 | Stage IIIC | Cutaneous | 4.2 | N/A (BRAF negative) | No |
| Chest Wall | Male | 59 | Stage IIIC | Cutaneous | 10 | BRAF V600E | No |
| Right Upper Arm | Male | 62 | Stage IIIC | Cutaneous | 2.5 | NRAS Q61K | No |
| Mid back skin | Male | 66 | Stage IIID | Cutaneous | 12 | no data | No |
| Right posterior thigh | Female | 67 | Stage IIIC | Cutaneous | 2.85 | BRAF V600E, ALK G667R, SMO V270I | No |
| Left foot, digits 2 and | Male | 74 | Stage IIIB | Acral lentiginous | 8 | no data | No |
| Left plantar forefoot | Female | 62 | Stage IIIC | Acral lentiginous | 6.8 | no data | No |
| Left Upper back | Male | 28 | Stage IIIB | Cutaneous | 2.2 | no data | No |
| Right leg | Male | 56 | Stage IIIC | Cutaneous | 6.9 | BRAF V600E | No |

#### Validation cohort

| Lymph node (LN) anatomic location | Stage at Excision | Type | Mutation status | Treatment before excision |
|---|---|---|---|---|
| Upper back skin and LN | IB | Unknown | BRAF Mut, NRAS WT, NF1 WT | No |
| Upper back skin and LN | IB | Unknown | BRAF Mut, NRAS WT, NF1 WT | No |
| Right axillary LN | IV | Unknown | BRAF WT, NRAS Mut, NF1 WT | No |
| Superficial inguinal LN | IIIB | Unclassified | BRAF WT, NRAS Mut, NF1 WT | No |
| Left axillary LN | IB | Superficial Spreading Melanom | BRAF Mut, NRAS WT, NF1 WT | No |
| Left axillary LN | IB | Superficial Spreading Melanom | BRAF Mut, NRAS WT, NF1 WT | No |
| Left inguinal LN | IIIC | Nodular melanoma | BRAF Mut, NRAS WT, NF1 WT | No |
| Intraparotid LN | IV | Unknown | BRAF WT, NRAS WT, NF1 Mut | No |
| Left axillary LN | IIIB | Unclassified | BRAF WT, NRAS WT, NF1 Mut | No |
| Left axillary LN | IIIB | Unclassified | BRAF WT, NRAS WT, NF1 Mut | No |
| Right supramohyoid LN | IB | Unclassified | BRAF WT, NRAS Mut, NF1 Mut | No |
| Right supramohyoid LN | IB | Unclassified | BRAF WT, NRAS Mut, NF1 Mut | No |
| Left axillary LN | IB | Unclassified | BRAF WT, NRAS Mut, NF1 Mut | No |
| Right superficial inguinal L | IIIC | Unclassified | BRAF Mut, NRAS WT, NF1 Mut | No |
| Right inguinal LN | IIIC | Superficial Spreading Melanom | BRAF Mut, NRAS WT, NF1 WT | No |
| Right inguinal LN | IIIC | Superficial Spreading Melanom | BRAF Mut, NRAS WT, NF1 WT | No |
| Left groin LN | II | Unclassified | BRAF WT, NRAS WT, NF1 WT | No |
| Left groin LN | II | Unclassified | BRAF WT, NRAS WT, NF1 WT | No |
