## Supplementary figures and images for "Lymphatic egress recycles tumor-experienced effector CD8 T cells to sustain immune surveillance"

### Supplemental Table 5

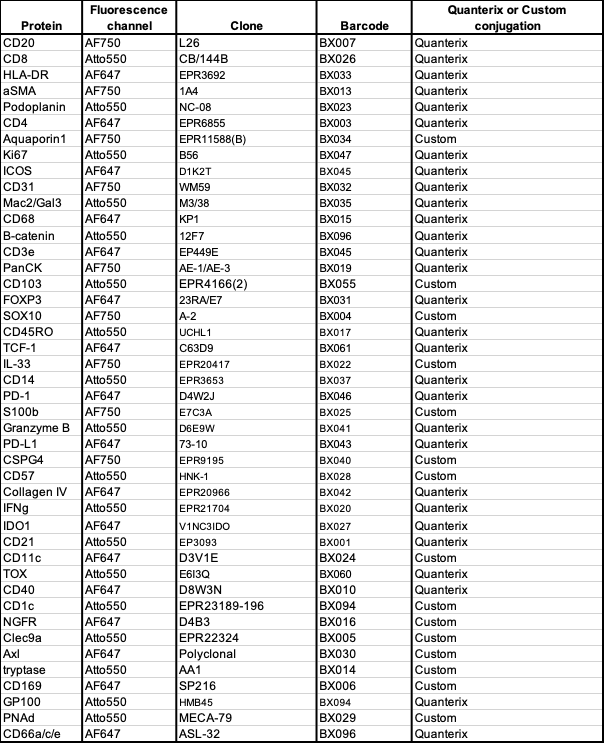
